## Supplementary material for "Distinct pathway for evolution of enhanced receptor binding and cell entry in SARS-like bat coronaviruses": PDB Validation Form

### Full wwPDB EM Validation Report ⓘ

Jun 18, 2024 – 10:55 AM EDT

| | | |
|---|---|---|
| PDB ID | : | 9CAS |
| EMDB ID | : | EMD-45402 |
| Title | : | Bat SARS-like Coronavirus RsSHC014 Spike Protein |
| Deposited on | : | 2024-06-17 |
| Resolution | : | 3.10 Å(reported) |

**This wwPDB validation report is for manuscript review**

This is a Full wwPDB EM Validation Report.

This report is produced by the wwPDB biocuration pipeline after annotation of the structure.

We welcome your comments at **
A user guide is available at
https://www.wwpdb.org/validation/2017/EMValidationReportHelp
with specific help available everywhere you see the ⓘ symbol.

The types of validation reports are described at
http://www.wwpdb.org/validation/2017/FAQs#types.

The following versions of software and data (see references ⓘ) were used in the production of this report:

| | | |
|---|---|---|
| EMDB validation analysis | : | 0.0.1.dev92 |
| MolProbity | : | 4.02b-467 |
| Percentile statistics | : | 20191225.v01 (using entries in the PDB archive December 25th 2019) |
| MapQ | : | 1.9.13 |
| Ideal geometry (proteins) | : | Engh & Huber (2001) |
| Ideal geometry (DNA, RNA) | : | Parkinson et al. (1996) |
| Validation Pipeline (wwPDB-VP) | : | 2.37.1 |

For Manuscript Review

### 1 Overall quality at a glance (i)

The following experimental techniques were used to determine the structure:
*ELECTRON MICROSCOPY*

The reported resolution of this entry is 3.10 Å.

Percentile scores (ranging between 0-100) for global validation metrics of the entry are shown in the following graphic. The table shows the number of entries on which the scores are based.

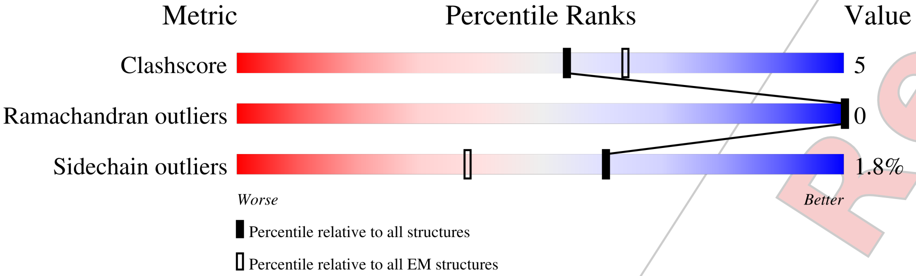


| Metric | Whole archive (#Entries) | EM structures (#Entries) |
|---|---|---|
| Clashscore | 158937 | 4297 |
| Ramachandran outliers | 154571 | 4023 |
| Sidechain outliers | 154315 | 3826 |

The table below summarises the geometric issues observed across the polymeric chains and their fit to the map. The red, orange, yellow and green segments of the bar indicate the fraction of residues that contain outliers for >=3, 2, 1 and 0 types of geometric quality criteria respectively. A grey segment represents the fraction of residues that are not modelled. The numeric value for each fraction is indicated below the corresponding segment, with a dot representing fractions <=5% The upper red bar (where present) indicates the fraction of residues that have poor fit to the EM map (all-atom inclusion < 40%). The numeric value is given above the bar.

| Mol | Chain | Length | Quality of chain |
|---|---|---|---|
| 1 | A | 1271 | 66% 11% 23% |
| 1 | B | 1271 | 67% 10% 23% |
| 1 | C | 1271 | 66% 11% 23% |

For Manuscript Review

#### 2 Entry composition (i)

There is only 1 type of molecule in this entry. The entry contains 45023 atoms, of which 22251 are hydrogens and 0 are deuteriums.

In the tables below, the AltConf column contains the number of residues with at least one atom in alternate conformation and the Trace column contains the number of residues modelled with at most 2 atoms.

- Molecule 1 is a protein called Spike glycoprotein.

| Mol | Chain | Residues | Atoms | | | | | | AltConf | Trace |
|---|---|---|---|---|---|---|---|---|---|---|
| 1 | A | 975 | Total 15009 | C 4847 | H 7419 | N 1270 | O 1433 | S 40 | 0 | 0 |
| 1 | B | 974 | Total 14999 | C 4845 | H 7413 | N 1269 | O 1432 | S 40 | 0 | 0 |
| 1 | C | 976 | Total 15015 | C 4850 | H 7419 | N 1271 | O 1435 | S 40 | 0 | 0 |

There are 240 discrepancies between the modelled and reference sequences:

| Chain | Residue | Modelled | Actual | Comment | Reference |
|---|---|---|---|---|---|
| A | 1192 | GLY | - | expression tag | UNP U5WLK5 |
| A | 1193 | SER | - | expression tag | UNP U5WLK5 |
| A | 1194 | GLY | - | expression tag | UNP U5WLK5 |
| A | 1195 | TYR | - | expression tag | UNP U5WLK5 |
| A | 1196 | ILE | - | expression tag | UNP U5WLK5 |
| A | 1197 | PRO | - | expression tag | UNP U5WLK5 |
| A | 1198 | GLU | - | expression tag | UNP U5WLK5 |
| A | 1199 | ALA | - | expression tag | UNP U5WLK5 |
| A | 1200 | PRO | - | expression tag | UNP U5WLK5 |
| A | 1201 | ARG | - | expression tag | UNP U5WLK5 |
| A | 1202 | ASP | - | expression tag | UNP U5WLK5 |
| A | 1203 | GLY | - | expression tag | UNP U5WLK5 |
| A | 1204 | GLN | - | expression tag | UNP U5WLK5 |
| A | 1205 | ALA | - | expression tag | UNP U5WLK5 |
| A | 1206 | TYR | - | expression tag | UNP U5WLK5 |
| A | 1207 | VAL | - | expression tag | UNP U5WLK5 |
| A | 1208 | ARG | - | expression tag | UNP U5WLK5 |
| A | 1209 | LYS | - | expression tag | UNP U5WLK5 |
| A | 1210 | ASP | - | expression tag | UNP U5WLK5 |
| A | 1211 | GLY | - | expression tag | UNP U5WLK5 |
| A | 1212 | GLU | - | expression tag | UNP U5WLK5 |
| A | 1213 | TRP | - | expression tag | UNP U5WLK5 |
| A | 1214 | VAL | - | expression tag | UNP U5WLK5 |
| A | 1215 | LEU | - | expression tag | UNP U5WLK5 |

Continued on next page...

For Manuscript Review

Continued from previous page...

| Chain | Residue | Modelled | Actual | Comment | Reference |
|---|---|---|---|---|---|
| A | 1216 | LEU | - | expression tag | UNP U5WLK5 |
| A | 1217 | SER | - | expression tag | UNP U5WLK5 |
| A | 1218 | THR | - | expression tag | UNP U5WLK5 |
| A | 1219 | PHE | - | expression tag | UNP U5WLK5 |
| A | 1220 | LEU | - | expression tag | UNP U5WLK5 |
| A | 1221 | GLY | - | expression tag | UNP U5WLK5 |
| A | 1222 | ARG | - | expression tag | UNP U5WLK5 |
| A | 1223 | SER | - | expression tag | UNP U5WLK5 |
| A | 1224 | LEU | - | expression tag | UNP U5WLK5 |
| A | 1225 | GLU | - | expression tag | UNP U5WLK5 |
| A | 1226 | VAL | - | expression tag | UNP U5WLK5 |
| A | 1227 | LEU | - | expression tag | UNP U5WLK5 |
| A | 1228 | PHE | - | expression tag | UNP U5WLK5 |
| A | 1229 | GLN | - | expression tag | UNP U5WLK5 |
| A | 1230 | GLY | - | expression tag | UNP U5WLK5 |
| A | 1231 | PRO | - | expression tag | UNP U5WLK5 |
| A | 1232 | GLY | - | expression tag | UNP U5WLK5 |
| A | 1233 | HIS | - | expression tag | UNP U5WLK5 |
| A | 1234 | HIS | - | expression tag | UNP U5WLK5 |
| A | 1235 | HIS | - | expression tag | UNP U5WLK5 |
| A | 1236 | HIS | - | expression tag | UNP U5WLK5 |
| A | 1237 | HIS | - | expression tag | UNP U5WLK5 |
| A | 1238 | HIS | - | expression tag | UNP U5WLK5 |
| A | 1239 | HIS | - | expression tag | UNP U5WLK5 |
| A | 1240 | HIS | - | expression tag | UNP U5WLK5 |
| A | 1241 | SER | - | expression tag | UNP U5WLK5 |
| A | 1242 | ALA | - | expression tag | UNP U5WLK5 |
| A | 1243 | TRP | - | expression tag | UNP U5WLK5 |
| A | 1244 | SER | - | expression tag | UNP U5WLK5 |
| A | 1245 | HIS | - | expression tag | UNP U5WLK5 |
| A | 1246 | PRO | - | expression tag | UNP U5WLK5 |
| A | 1247 | GLN | - | expression tag | UNP U5WLK5 |
| A | 1248 | PHE | - | expression tag | UNP U5WLK5 |
| A | 1249 | GLU | - | expression tag | UNP U5WLK5 |
| A | 1250 | LYS | - | expression tag | UNP U5WLK5 |
| A | 1251 | GLY | - | expression tag | UNP U5WLK5 |
| A | 1252 | GLY | - | expression tag | UNP U5WLK5 |
| A | 1253 | GLY | - | expression tag | UNP U5WLK5 |
| A | 1254 | SER | - | expression tag | UNP U5WLK5 |
| A | 1255 | GLY | - | expression tag | UNP U5WLK5 |
| A | 1256 | GLY | - | expression tag | UNP U5WLK5 |
| A | 1257 | GLY | - | expression tag | UNP U5WLK5 |

Continued on next page...

For Manuscript Review

Continued from previous page...

| Chain | Residue | Modelled | Actual | Comment | Reference |
|---|---|---|---|---|---|
| A | 1258 | GLY | - | expression tag | UNP U5WLK5 |
| A | 1259 | SER | - | expression tag | UNP U5WLK5 |
| A | 1260 | GLY | - | expression tag | UNP U5WLK5 |
| A | 1261 | GLY | - | expression tag | UNP U5WLK5 |
| A | 1262 | SER | - | expression tag | UNP U5WLK5 |
| A | 1263 | ALA | - | expression tag | UNP U5WLK5 |
| A | 1264 | TRP | - | expression tag | UNP U5WLK5 |
| A | 1265 | SER | - | expression tag | UNP U5WLK5 |
| A | 1266 | HIS | - | expression tag | UNP U5WLK5 |
| A | 1267 | PRO | - | expression tag | UNP U5WLK5 |
| A | 1268 | GLN | - | expression tag | UNP U5WLK5 |
| A | 1269 | PHE | - | expression tag | UNP U5WLK5 |
| A | 1270 | GLU | - | expression tag | UNP U5WLK5 |
| A | 1271 | LYS | - | expression tag | UNP U5WLK5 |
| B | 1192 | GLY | - | expression tag | UNP U5WLK5 |
| B | 1193 | SER | - | expression tag | UNP U5WLK5 |
| B | 1194 | GLY | - | expression tag | UNP U5WLK5 |
| B | 1195 | TYR | - | expression tag | UNP U5WLK5 |
| B | 1196 | ILE | - | expression tag | UNP U5WLK5 |
| B | 1197 | PRO | - | expression tag | UNP U5WLK5 |
| B | 1198 | GLU | - | expression tag | UNP U5WLK5 |
| B | 1199 | ALA | - | expression tag | UNP U5WLK5 |
| B | 1200 | PRO | - | expression tag | UNP U5WLK5 |
| B | 1201 | ARG | - | expression tag | UNP U5WLK5 |
| B | 1202 | ASP | - | expression tag | UNP U5WLK5 |
| B | 1203 | GLY | - | expression tag | UNP U5WLK5 |
| B | 1204 | GLN | - | expression tag | UNP U5WLK5 |
| B | 1205 | ALA | - | expression tag | UNP U5WLK5 |
| B | 1206 | TYR | - | expression tag | UNP U5WLK5 |
| B | 1207 | VAL | - | expression tag | UNP U5WLK5 |
| B | 1208 | ARG | - | expression tag | UNP U5WLK5 |
| B | 1209 | LYS | - | expression tag | UNP U5WLK5 |
| B | 1210 | ASP | - | expression tag | UNP U5WLK5 |
| B | 1211 | GLY | - | expression tag | UNP U5WLK5 |
| B | 1212 | GLU | - | expression tag | UNP U5WLK5 |
| B | 1213 | TRP | - | expression tag | UNP U5WLK5 |
| B | 1214 | VAL | - | expression tag | UNP U5WLK5 |
| B | 1215 | LEU | - | expression tag | UNP U5WLK5 |
| B | 1216 | LEU | - | expression tag | UNP U5WLK5 |
| B | 1217 | SER | - | expression tag | UNP U5WLK5 |
| B | 1218 | THR | - | expression tag | UNP U5WLK5 |
| B | 1219 | PHE | - | expression tag | UNP U5WLK5 |

Continued on next page...

For Manuscript Review

*Continued from previous page...*

| Chain | Residue | Modelled | Actual | Comment | Reference |
|---|---|---|---|---|---|
| B | 1220 | LEU | - | expression tag | UNP U5WLK5 |
| B | 1221 | GLY | - | expression tag | UNP U5WLK5 |
| B | 1222 | ARG | - | expression tag | UNP U5WLK5 |
| B | 1223 | SER | - | expression tag | UNP U5WLK5 |
| B | 1224 | LEU | - | expression tag | UNP U5WLK5 |
| B | 1225 | GLU | - | expression tag | UNP U5WLK5 |
| B | 1226 | VAL | - | expression tag | UNP U5WLK5 |
| B | 1227 | LEU | - | expression tag | UNP U5WLK5 |
| B | 1228 | PHE | - | expression tag | UNP U5WLK5 |
| B | 1229 | GLN | - | expression tag | UNP U5WLK5 |
| B | 1230 | GLY | - | expression tag | UNP U5WLK5 |
| B | 1231 | PRO | - | expression tag | UNP U5WLK5 |
| B | 1232 | GLY | - | expression tag | UNP U5WLK5 |
| B | 1233 | HIS | - | expression tag | UNP U5WLK5 |
| B | 1234 | HIS | - | expression tag | UNP U5WLK5 |
| B | 1235 | HIS | - | expression tag | UNP U5WLK5 |
| B | 1236 | HIS | - | expression tag | UNP U5WLK5 |
| B | 1237 | HIS | - | expression tag | UNP U5WLK5 |
| B | 1238 | HIS | - | expression tag | UNP U5WLK5 |
| B | 1239 | HIS | - | expression tag | UNP U5WLK5 |
| B | 1240 | HIS | - | expression tag | UNP U5WLK5 |
| B | 1241 | SER | - | expression tag | UNP U5WLK5 |
| B | 1242 | ALA | - | expression tag | UNP U5WLK5 |
| B | 1243 | TRP | - | expression tag | UNP U5WLK5 |
| B | 1244 | SER | - | expression tag | UNP U5WLK5 |
| B | 1245 | HIS | - | expression tag | UNP U5WLK5 |
| B | 1246 | PRO | - | expression tag | UNP U5WLK5 |
| B | 1247 | GLN | - | expression tag | UNP U5WLK5 |
| B | 1248 | PHE | - | expression tag | UNP U5WLK5 |
| B | 1249 | GLU | - | expression tag | UNP U5WLK5 |
| B | 1250 | LYS | - | expression tag | UNP U5WLK5 |
| B | 1251 | GLY | - | expression tag | UNP U5WLK5 |
| B | 1252 | GLY | - | expression tag | UNP U5WLK5 |
| B | 1253 | GLY | - | expression tag | UNP U5WLK5 |
| B | 1254 | SER | - | expression tag | UNP U5WLK5 |
| B | 1255 | GLY | - | expression tag | UNP U5WLK5 |
| B | 1256 | GLY | - | expression tag | UNP U5WLK5 |
| B | 1257 | GLY | - | expression tag | UNP U5WLK5 |
| B | 1258 | GLY | - | expression tag | UNP U5WLK5 |
| B | 1259 | SER | - | expression tag | UNP U5WLK5 |
| B | 1260 | GLY | - | expression tag | UNP U5WLK5 |
| B | 1261 | GLY | - | expression tag | UNP U5WLK5 |

*Continued on next page...*

For Manuscript Review

Continued from previous page...

| Chain | Residue | Modelled | Actual | Comment | Reference |
|---|---|---|---|---|---|
| B | 1262 | SER | - | expression tag | UNP U5WLK5 |
| B | 1263 | ALA | - | expression tag | UNP U5WLK5 |
| B | 1264 | TRP | - | expression tag | UNP U5WLK5 |
| B | 1265 | SER | - | expression tag | UNP U5WLK5 |
| B | 1266 | HIS | - | expression tag | UNP U5WLK5 |
| B | 1267 | PRO | - | expression tag | UNP U5WLK5 |
| B | 1268 | GLN | - | expression tag | UNP U5WLK5 |
| B | 1269 | PHE | - | expression tag | UNP U5WLK5 |
| B | 1270 | GLU | - | expression tag | UNP U5WLK5 |
| B | 1271 | LYS | - | expression tag | UNP U5WLK5 |
| C | 1192 | GLY | - | expression tag | UNP U5WLK5 |
| C | 1193 | SER | - | expression tag | UNP U5WLK5 |
| C | 1194 | GLY | - | expression tag | UNP U5WLK5 |
| C | 1195 | TYR | - | expression tag | UNP U5WLK5 |
| C | 1196 | ILE | - | expression tag | UNP U5WLK5 |
| C | 1197 | PRO | - | expression tag | UNP U5WLK5 |
| C | 1198 | GLU | - | expression tag | UNP U5WLK5 |
| C | 1199 | ALA | - | expression tag | UNP U5WLK5 |
| C | 1200 | PRO | - | expression tag | UNP U5WLK5 |
| C | 1201 | ARG | - | expression tag | UNP U5WLK5 |
| C | 1202 | ASP | - | expression tag | UNP U5WLK5 |
| C | 1203 | GLY | - | expression tag | UNP U5WLK5 |
| C | 1204 | GLN | - | expression tag | UNP U5WLK5 |
| C | 1205 | ALA | - | expression tag | UNP U5WLK5 |
| C | 1206 | TYR | - | expression tag | UNP U5WLK5 |
| C | 1207 | VAL | - | expression tag | UNP U5WLK5 |
| C | 1208 | ARG | - | expression tag | UNP U5WLK5 |
| C | 1209 | LYS | - | expression tag | UNP U5WLK5 |
| C | 1210 | ASP | - | expression tag | UNP U5WLK5 |
| C | 1211 | GLY | - | expression tag | UNP U5WLK5 |
| C | 1212 | GLU | - | expression tag | UNP U5WLK5 |
| C | 1213 | TRP | - | expression tag | UNP U5WLK5 |
| C | 1214 | VAL | - | expression tag | UNP U5WLK5 |
| C | 1215 | LEU | - | expression tag | UNP U5WLK5 |
| C | 1216 | LEU | - | expression tag | UNP U5WLK5 |
| C | 1217 | SER | - | expression tag | UNP U5WLK5 |
| C | 1218 | THR | - | expression tag | UNP U5WLK5 |
| C | 1219 | PHE | - | expression tag | UNP U5WLK5 |
| C | 1220 | LEU | - | expression tag | UNP U5WLK5 |
| C | 1221 | GLY | - | expression tag | UNP U5WLK5 |
| C | 1222 | ARG | - | expression tag | UNP U5WLK5 |
| C | 1223 | SER | - | expression tag | UNP U5WLK5 |

Continued on next page...

For Manuscript Review

*Continued from previous page...*

| Chain | Residue | Modelled | Actual | Comment | Reference |
|---|---|---|---|---|---|
| C | 1224 | LEU | - | expression tag | UNP U5WLK5 |
| C | 1225 | GLU | - | expression tag | UNP U5WLK5 |
| C | 1226 | VAL | - | expression tag | UNP U5WLK5 |
| C | 1227 | LEU | - | expression tag | UNP U5WLK5 |
| C | 1228 | PHE | - | expression tag | UNP U5WLK5 |
| C | 1229 | GLN | - | expression tag | UNP U5WLK5 |
| C | 1230 | GLY | - | expression tag | UNP U5WLK5 |
| C | 1231 | PRO | - | expression tag | UNP U5WLK5 |
| C | 1232 | GLY | - | expression tag | UNP U5WLK5 |
| C | 1233 | HIS | - | expression tag | UNP U5WLK5 |
| C | 1234 | HIS | - | expression tag | UNP U5WLK5 |
| C | 1235 | HIS | - | expression tag | UNP U5WLK5 |
| C | 1236 | HIS | - | expression tag | UNP U5WLK5 |
| C | 1237 | HIS | - | expression tag | UNP U5WLK5 |
| C | 1238 | HIS | - | expression tag | UNP U5WLK5 |
| C | 1239 | HIS | - | expression tag | UNP U5WLK5 |
| C | 1240 | HIS | - | expression tag | UNP U5WLK5 |
| C | 1241 | SER | - | expression tag | UNP U5WLK5 |
| C | 1242 | ALA | - | expression tag | UNP U5WLK5 |
| C | 1243 | TRP | - | expression tag | UNP U5WLK5 |
| C | 1244 | SER | - | expression tag | UNP U5WLK5 |
| C | 1245 | HIS | - | expression tag | UNP U5WLK5 |
| C | 1246 | PRO | - | expression tag | UNP U5WLK5 |
| C | 1247 | GLN | - | expression tag | UNP U5WLK5 |
| C | 1248 | PHE | - | expression tag | UNP U5WLK5 |
| C | 1249 | GLU | - | expression tag | UNP U5WLK5 |
| C | 1250 | LYS | - | expression tag | UNP U5WLK5 |
| C | 1251 | GLY | - | expression tag | UNP U5WLK5 |
| C | 1252 | GLY | - | expression tag | UNP U5WLK5 |
| C | 1253 | GLY | - | expression tag | UNP U5WLK5 |
| C | 1254 | SER | - | expression tag | UNP U5WLK5 |
| C | 1255 | GLY | - | expression tag | UNP U5WLK5 |
| C | 1256 | GLY | - | expression tag | UNP U5WLK5 |
| C | 1257 | GLY | - | expression tag | UNP U5WLK5 |
| C | 1258 | GLY | - | expression tag | UNP U5WLK5 |
| C | 1259 | SER | - | expression tag | UNP U5WLK5 |
| C | 1260 | GLY | - | expression tag | UNP U5WLK5 |
| C | 1261 | GLY | - | expression tag | UNP U5WLK5 |
| C | 1262 | SER | - | expression tag | UNP U5WLK5 |
| C | 1263 | ALA | - | expression tag | UNP U5WLK5 |
| C | 1264 | TRP | - | expression tag | UNP U5WLK5 |
| C | 1265 | SER | - | expression tag | UNP U5WLK5 |

*Continued on next page...*

For Manuscript Review

*Continued from previous page...*

| Chain | Residue | Modelled | Actual | Comment | Reference |
|---|---|---|---|---|---|
| C | 1266 | HIS | - | expression tag | UNP U5WLK5 |
| C | 1267 | PRO | - | expression tag | UNP U5WLK5 |
| C | 1268 | GLN | - | expression tag | UNP U5WLK5 |
| C | 1269 | PHE | - | expression tag | UNP U5WLK5 |
| C | 1270 | GLU | - | expression tag | UNP U5WLK5 |
| C | 1271 | LYS | - | expression tag | UNP U5WLK5 |

For Manuscript Review

### 3 Residue-property plots ⓘ

These plots are drawn for all protein, RNA, DNA and oligosaccharide chains in the entry. The first graphic for a chain summarises the proportions of the various outlier classes displayed in the second graphic. The second graphic shows the sequence view annotated by issues in geometry and atom inclusion in map density. Residues are color-coded according to the number of geometric quality criteria for which they contain at least one outlier: green = 0, yellow = 1, orange = 2 and red = 3 or more. A red diamond above a residue indicates a poor fit to the EM map for this residue (all-atom inclusion < 40%). Stretches of 2 or more consecutive residues without any outlier are shown as a green connector. Residues present in the sample, but not in the model, are shown in grey.

- Molecule 1: Spike glycoprotein

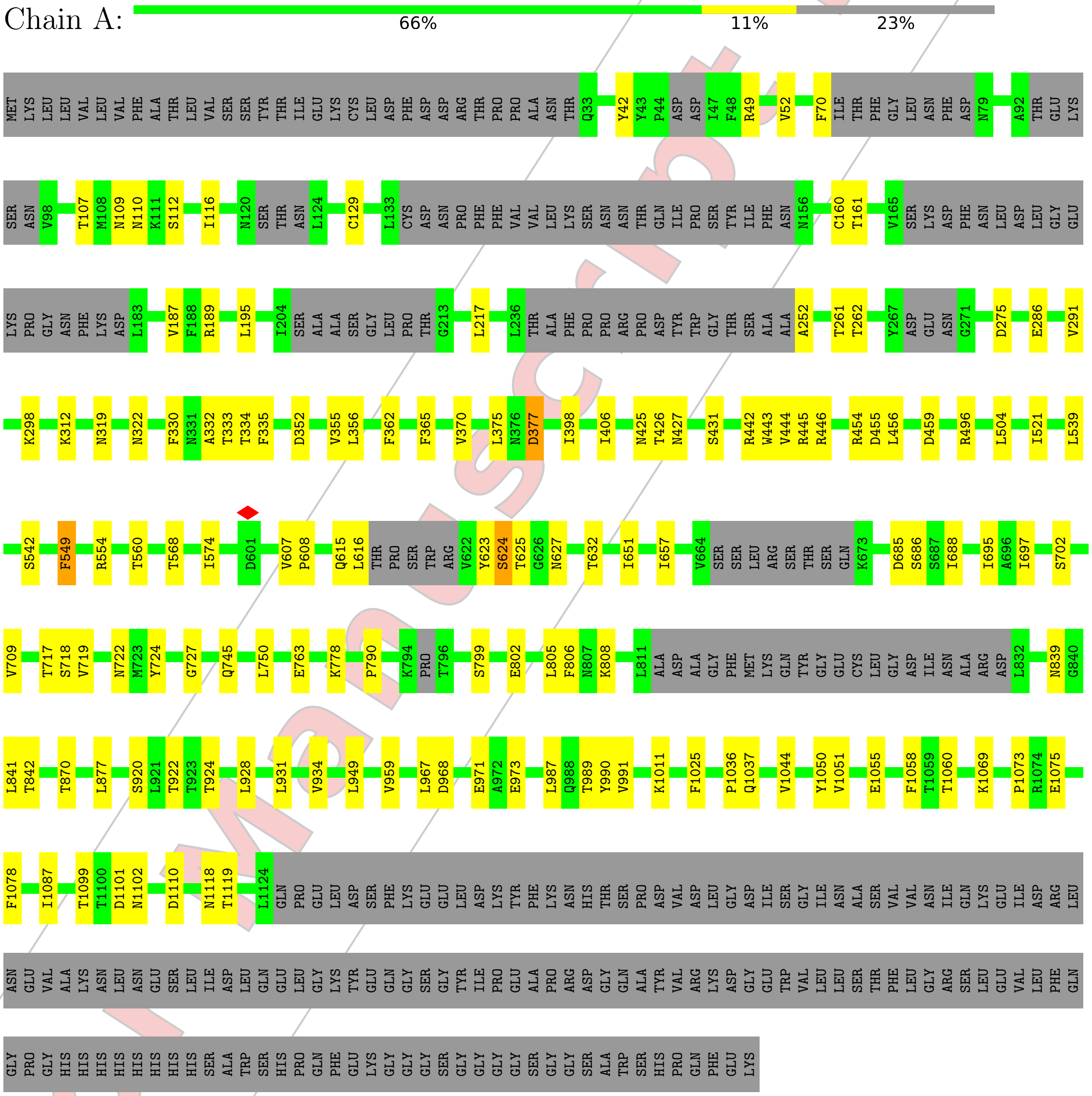


- Molecule 1: Spike glycoprotein

For Manuscript Review

Chain B: 67% 10% 23%

MET LYS LEU LEU VAL LEU VAL PHE ALA THR LEU VAL SER SER TYR THR ILE GLU LYS CYS LEU ASP PHE ASP ASP ARG THR PRO PRO ALA ASN THR Q33 P44 ASP ASP I47 F70 ILE THR PHE GLY LEU ASN PHE ASP N79 A92 THR GLU LYS SER ASN V98 N109 N110

N120 SER THR ASN L124 L133 CYS ASP ASN PRO PHE PHE VAL VAL LEU LYS SER ASN ASN THR GLN ILE PRO SER TYR ILE PHE ASN N156 V165 SER LYS ASP PHE ASN LEU ASP LEU GLY GLU LYS PRO GLY ASN PHE LYS ASP L183 V187 F188 R189 L195 I204 SER

ALA ALA SER GLY LEU PRO THR GLY F214 L217 L236 THR ALA PHE PRO PRO ARG PRO ASP TYR TRP GLY THR SER ALA ALA A252 T261 T262 L265 K266 Y267 ASP GLU ASN G271 T272 I273 S280 E286 V291 K298 N305 E313 V314 V315 N319 F335

D352 V355 L356 F362 F365 S371 I398 I406 N425 T426 N427 S431 R442 W443 V444 R445 R446 Y452 E453 R454 D455 S462 S501 L516 S517 T518 D519 L539 R554 T560 V563 R564 T568 I574 S578 F579 Q600

V607 P608 L616 THR PRO SER TRP ARG V622 Y623 S624 N627 N628 T632 I651 G656 I657 V664 SER SER LEU ARG SER THR SER GLN K673 L682 D685 S686 S687 I688 I695 A696 I697 S702 V709 S718 V719 N722 M723 Y724 G727 Q745

E763 K778 P790 K794 PRO T796 S799 E802 L805 F806 N807 K808 L811 ALA ASP ALA GLY PHE MET LYS GLN TYR GLY GLU CYS LEU GLY ASP ILE ASN ALA ARG ASP L832 N839 G840 L841 T842 T870 L877 A913 I917 T922 T923 T924 L928

L931 V934 L949 V959 L960 R966 L967 D968 E971 A972 E973 I976 L987 Q988 T989 Q993 K1011 F1025 P1036 Q1037 V1044 V1051 E1055 F1058 E1075 T1099 T1100 D1101 N1102 D1110 N1118 T1119 V1120 Y1121 L1124 GLN PRO GLU LEU

ASP SER PHE LYS GLU GLU LEU ASP LYS TYR PHE LYS ASN HIS THR SER PRO ASP VAL ASP LEU GLY ASP ILE SER GLY ILE ASN ALA SER VAL VAL ASN ILE GLN LYS GLU ILE ASP ARG LEU ASN GLU VAL ALA LYS ASN LEU ASN GLU SER LEU ILE ASP LEU GLN GLU LEU GLY LYS

TYR GLU GLN GLY SER GLY TYR ILE PRO GLU ALA PRO ARG ASP GLY GLN ALA TYR VAL ARG LYS ASP GLY GLU TRP VAL LEU LEU SER THR PHE LEU GLY ARG SER LEU GLU VAL LEU PHE GLN GLY PRO GLY HIS HIS HIS HIS HIS HIS HIS HIS SER ALA TRP SER HIS PRO GLN PHE

GLU LYS GLY GLY GLY SER GLY GLY GLY SER GLY GLY SER ALA TRP SER HIS PRO GLN PHE GLU LYS

- Molecule 1: Spike glycoprotein

Chain C: 66% 11% 23%

MET LYS LEU LEU VAL LEU VAL PHE ALA THR LEU VAL SER SER TYR THR ILE GLU LYS CYS LEU ASP PHE ASP ASP ARG THR PRO PRO ALA ASN THR Q33 R39 P44 ASP ASP I47 F48 R49 S50 N51 V52 F70 ILE THR PHE GLY LEU ASN PHE ASP N79 A92 THR GLU

LYS SER ASN V98 T107 M108 N109 N110 K111 S112 I116 N120 SER THR ASN L124 A128 C129 L133 CYS ASP ASN PRO PHE PHE VAL VAL LEU LYS SER ASN ASN THR GLN ILE PRO SER TYR ILE PHE ASN N156 C160 T161 S166 LYS ASP PHE ASN LEU ASP LEU GLY

GLU LYS PRO GLY ASN PHE LYS ASP L183 R184 E185 F186 V187 F188 R189 L195 I204 SER ALA ALA SER GLY LEU PRO THR G213 L217 L236 THR ALA PHE PRO PRO ARG PRO ASP TYR TRP GLY THR SER ALA ALA A252 T261 T262 Y267 ASP GLU ASN G271 D275

E286 V291 K298 E313 V314 V315 N319 N322 F330 N348 C349 V350 A351 D352 V355 L356 F362 F365 S371 D377 N425 T426 N427 S431 R442 R445 R446 Y452 E453 R454 D455 S501 L539 S542 R545 R554 T560

For Manuscript Review

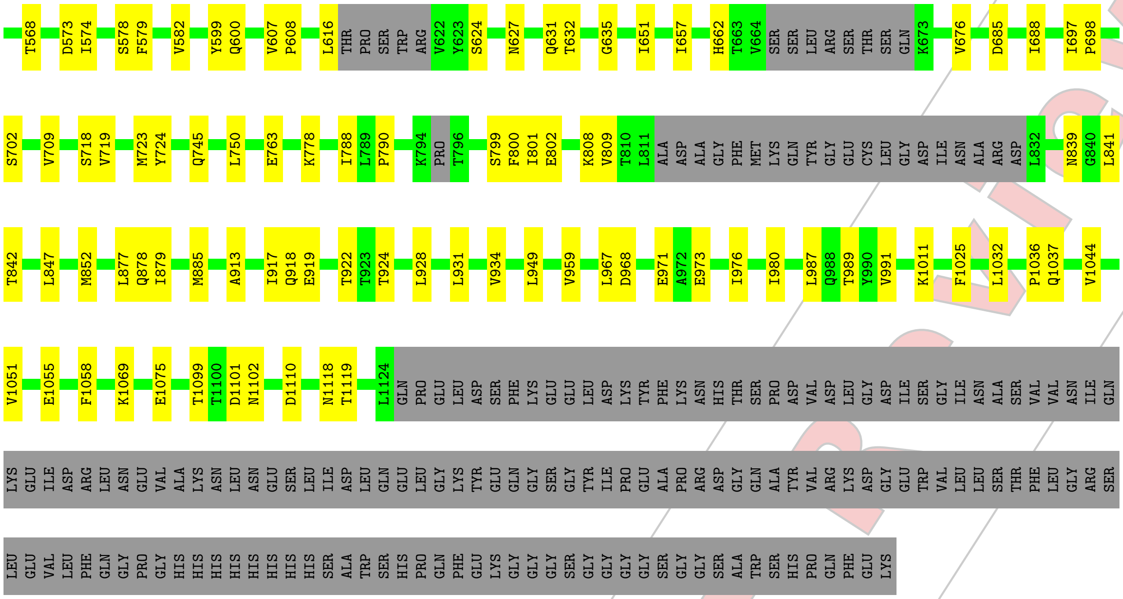


For Manuscript Review

### 4 Experimental information (i)

| Property | Value | Source |
|---|---|---|
| EM reconstruction method | SINGLE PARTICLE | Depositor |
| Imposed symmetry | POINT, Not provided | |
| Number of particles used | 76442 | Depositor |
| Resolution determination method | FSC 0.143 CUT-OFF | Depositor |
| CTF correction method | PHASE FLIPPING AND AMPLITUDE CORRECTION | Depositor |
| Microscope | TFS GLACIOS | Depositor |
| Voltage (kV) | 200 | Depositor |
| Electron dose $(e^{-}/\text{Å}^2)$ | 50 | Depositor |
| Minimum defocus (nm) | 1000 | Depositor |
| Maximum defocus (nm) | 2000 | Depositor |
| Magnification | Not provided | |
| Image detector | FEI FALCON IV (4k x 4k) | Depositor |
| Maximum map value | 1.458 | Depositor |
| Minimum map value | -1.044 | Depositor |
| Average map value | -0.000 | Depositor |
| Map value standard deviation | 0.026 | Depositor |
| Recommended contour level | 0.18 | Depositor |
| Map size (Å) | 481.28, 481.28, 481.28 | wwPDB |
| Map dimensions | 512, 512, 512 | wwPDB |
| Map angles (°) | 90.0, 90.0, 90.0 | wwPDB |
| Pixel spacing (Å) | 0.94, 0.94, 0.94 | Depositor |

For Manuscript Review

### 5 Model quality (i)

#### 5.1 Standard geometry (i)

The Z score for a bond length (or angle) is the number of standard deviations the observed value is removed from the expected value. A bond length (or angle) with $|Z| > 5$ is considered an outlier worth inspection. RMSZ is the root-mean-square of all Z scores of the bond lengths (or angles).

| Mol | Chain | Bond lengths | | Bond angles | |
|---|---|---|---|---|---|
| | | RMSZ | $\#|Z| > 5$ | RMSZ | $\#|Z| > 5$ |
| 1 | A | 0.28 | 0/7753 | 0.49 | 0/10532 |
| 1 | B | 0.28 | 0/7749 | 0.49 | 0/10527 |
| 1 | C | 0.28 | 0/7759 | 0.49 | 0/10540 |
| All | All | 0.28 | 0/23261 | 0.49 | 0/31599 |

There are no bond length outliers.

There are no bond angle outliers.

There are no chirality outliers.

There are no planarity outliers.

#### 5.2 Too-close contacts (i)

In the following table, the Non-H and H(model) columns list the number of non-hydrogen atoms and hydrogen atoms in the chain respectively. The H(added) column lists the number of hydrogen atoms added and optimized by MolProbity. The Clashes column lists the number of clashes within the asymmetric unit, whereas Symm-Clashes lists symmetry-related clashes.

| Mol | Chain | Non-H | H(model) | H(added) | Clashes | Symm-Clashes |
|---|---|---|---|---|---|---|
| 1 | A | 7590 | 7419 | 7426 | 92 | 0 |
| 1 | B | 7586 | 7413 | 7423 | 83 | 0 |
| 1 | C | 7596 | 7419 | 7431 | 93 | 0 |
| All | All | 22772 | 22251 | 22280 | 244 | 0 |

The all-atom clashscore is defined as the number of clashes found per 1000 atoms (including hydrogen atoms). The all-atom clashscore for this structure is 5.

All (244) close contacts within the same asymmetric unit are listed below, sorted by their clash magnitude.

| Atom-1 | Atom-2 | Interatomic distance (Å) | Clash overlap (Å) |
|---|---|---|---|
| 1:A:70:PHE:O | 1:A:252:ALA:N | 2.07 | 0.86 |

*Continued on next page...*

For Manuscript Review

*Continued from previous page...*

| Atom-1 | Atom-2 | Interatomic distance (Å) | Clash overlap (Å) |
|---|---|---|---|
| 1:A:427:ASN:O | 1:A:431:SER:OG | 1.95 | 0.83 |
| 1:C:427:ASN:O | 1:C:431:SER:OG | 1.97 | 0.82 |
| 1:B:427:ASN:O | 1:B:431:SER:OG | 1.98 | 0.80 |
| 1:C:49:ARG:O | 1:C:271:GLY:N | 2.17 | 0.78 |
| 1:A:333:THR:HG23 | 1:A:334:THR:HG23 | 1.67 | 0.76 |
| 1:A:778:LYS:NZ | 1:A:790:PRO:O | 2.19 | 0.74 |
| 1:B:709:VAL:HG12 | 1:B:1044:VAL:HG22 | 1.71 | 0.73 |
| 1:B:1011:LYS:NZ | 1:B:1025:PHE:O | 2.23 | 0.72 |
| 1:A:442:ARG:NH2 | 1:A:455:ASP:O | 2.24 | 0.71 |
| 1:C:107:THR:HG1 | 1:C:112:SER:HG | 1.26 | 0.71 |
| 1:C:539:LEU:HD23 | 1:C:574:ILE:HG22 | 1.73 | 0.70 |
| 1:C:778:LYS:NZ | 1:C:790:PRO:O | 2.24 | 0.70 |
| 1:B:442:ARG:NH2 | 1:B:455:ASP:O | 2.26 | 0.69 |
| 1:A:709:VAL:HG12 | 1:A:1044:VAL:HG22 | 1.74 | 0.69 |
| 1:A:129:CYS:HA | 1:A:160:CYS:HB2 | 1.75 | 0.68 |
| 1:C:709:VAL:HG12 | 1:C:1044:VAL:HG22 | 1.74 | 0.68 |
| 1:B:718:SER:OG | 1:B:842:THR:OG1 | 2.12 | 0.68 |
| 1:B:778:LYS:NZ | 1:B:790:PRO:O | 2.27 | 0.68 |
| 1:B:554:ARG:NH1 | 1:C:959:VAL:HG21 | 2.10 | 0.67 |
| 1:B:1075:GLU:OE1 | 1:B:1075:GLU:N | 2.28 | 0.66 |
| 1:A:959:VAL:HG21 | 1:C:554:ARG:NH1 | 2.11 | 0.65 |
| 1:C:718:SER:OG | 1:C:842:THR:OG1 | 2.15 | 0.64 |
| 1:C:49:ARG:HD3 | 1:C:52:VAL:HG11 | 1.79 | 0.64 |
| 1:B:1099:THR:OG1 | 1:B:1101:ASP:OD1 | 2.16 | 0.64 |
| 1:B:1110:ASP:OD1 | 1:B:1110:ASP:O | 2.15 | 0.64 |
| 1:A:1036:PRO:O | 1:A:1037:GLN:NE2 | 2.31 | 0.64 |
| 1:B:1055:GLU:HG2 | 1:C:877:LEU:HD22 | 1.80 | 0.64 |
| 1:A:107:THR:OG1 | 1:A:112:SER:OG | 2.10 | 0.64 |
| 1:A:1099:THR:OG1 | 1:A:1101:ASP:OD1 | 2.16 | 0.64 |
| 1:A:717:THR:O | 1:A:750:LEU:HD12 | 1.98 | 0.63 |
| 1:C:1075:GLU:N | 1:C:1075:GLU:OE1 | 2.30 | 0.63 |
| 1:A:554:ARG:HH12 | 1:B:959:VAL:HG21 | 1.63 | 0.62 |
| 1:C:697:ILE:HD11 | 1:C:1058:PHE:CE2 | 2.34 | 0.62 |
| 1:C:719:VAL:HG13 | 1:C:841:LEU:HD23 | 1.81 | 0.62 |
| 1:C:709:VAL:HG11 | 1:C:928:LEU:HD23 | 1.81 | 0.62 |
| 1:B:554:ARG:HH12 | 1:C:959:VAL:HG21 | 1.66 | 0.61 |
| 1:C:719:VAL:HG11 | 1:C:987:LEU:HD21 | 1.83 | 0.60 |
| 1:A:724:TYR:CE1 | 1:A:949:LEU:HD11 | 2.36 | 0.60 |
| 1:A:554:ARG:NH1 | 1:B:959:VAL:HG21 | 2.15 | 0.60 |
| 1:C:967:LEU:HD13 | 1:C:971:GLU:OE1 | 2.02 | 0.60 |
| 1:B:578:SER:O | 1:B:579:PHE:CD2 | 2.55 | 0.59 |

*Continued on next page...*

For Manuscript Review

*Continued from previous page...*

| Atom-1 | Atom-2 | Interatomic distance (Å) | Clash overlap (Å) |
|---|---|---|---|
| 1:C:442:ARG:NH2 | 1:C:455:ASP:O | 2.35 | 0.59 |
| 1:B:443:TRP:CE3 | 1:B:444:VAL:HG13 | 2.37 | 0.59 |
| 1:A:560:THR:O | 1:A:574:ILE:HD11 | 2.03 | 0.58 |
| 1:B:616:LEU:O | 1:B:616:LEU:HD23 | 2.04 | 0.58 |
| 1:B:452:TYR:OH | 1:B:501:SER:OG | 2.22 | 0.58 |
| 1:A:49:ARG:HD3 | 1:A:52:VAL:HG11 | 1.85 | 0.58 |
| 1:A:425:ASN:OD1 | 1:A:427:ASN:N | 2.34 | 0.58 |
| 1:C:616:LEU:HD23 | 1:C:616:LEU:O | 2.04 | 0.58 |
| 1:C:702:SER:OG | 1:C:1051:VAL:HG23 | 2.04 | 0.58 |
| 1:A:959:VAL:HG21 | 1:C:554:ARG:HH12 | 1.68 | 0.57 |
| 1:B:724:TYR:CE1 | 1:B:949:LEU:HD11 | 2.38 | 0.57 |
| 1:B:1036:PRO:O | 1:B:1037:GLN:NE2 | 2.37 | 0.57 |
| 1:C:724:TYR:CE1 | 1:C:949:LEU:HD11 | 2.39 | 0.57 |
| 1:C:685:ASP:N | 1:C:685:ASP:OD1 | 2.38 | 0.57 |
| 1:A:719:VAL:HG13 | 1:A:841:LEU:HD23 | 1.87 | 0.57 |
| 1:B:454:ARG:NH1 | 1:C:161:THR:HG21 | 2.19 | 0.57 |
| 1:C:1118:ASN:OD1 | 1:C:1119:THR:N | 2.37 | 0.56 |
| 1:B:280:SER:O | 1:B:280:SER:OG | 2.22 | 0.56 |
| 1:B:967:LEU:HD13 | 1:B:971:GLU:OE1 | 2.05 | 0.56 |
| 1:C:129:CYS:HA | 1:C:160:CYS:HB2 | 1.88 | 0.56 |
| 1:A:718:SER:OG | 1:A:842:THR:OG1 | 2.22 | 0.56 |
| 1:B:805:LEU:HD22 | 1:B:928:LEU:HD21 | 1.88 | 0.56 |
| 1:A:189:ARG:O | 1:A:195:LEU:HD12 | 2.06 | 0.56 |
| 1:A:799:SER:OG | 1:A:802:GLU:OE1 | 2.22 | 0.56 |
| 1:B:719:VAL:HG13 | 1:B:841:LEU:HD23 | 1.87 | 0.55 |
| 1:C:1099:THR:OG1 | 1:C:1101:ASP:OD1 | 2.24 | 0.55 |
| 1:B:454:ARG:HH11 | 1:C:161:THR:HG21 | 1.71 | 0.55 |
| 1:C:1036:PRO:O | 1:C:1037:GLN:NE2 | 2.39 | 0.55 |
| 1:A:702:SER:OG | 1:A:1051:VAL:HG23 | 2.06 | 0.55 |
| 1:C:452:TYR:OH | 1:C:501:SER:OG | 2.25 | 0.55 |
| 1:B:702:SER:OG | 1:B:1051:VAL:HG23 | 2.08 | 0.54 |
| 1:C:1110:ASP:OD1 | 1:C:1110:ASP:O | 2.25 | 0.54 |
| 1:B:286:GLU:OE1 | 1:B:616:LEU:HD21 | 2.07 | 0.54 |
| 1:C:425:ASN:OD1 | 1:C:427:ASN:N | 2.38 | 0.53 |
| 1:B:313:GLU:HG2 | 1:B:315:VAL:HG23 | 1.91 | 0.53 |
| 1:A:454:ARG:HD2 | 1:A:456:LEU:HD11 | 1.90 | 0.53 |
| 1:A:1118:ASN:OD1 | 1:A:1119:THR:N | 2.41 | 0.53 |
| 1:B:719:VAL:HG11 | 1:B:987:LEU:HD21 | 1.89 | 0.53 |
| 1:C:189:ARG:O | 1:C:195:LEU:HD12 | 2.09 | 0.53 |
| 1:C:1011:LYS:NZ | 1:C:1025:PHE:O | 2.42 | 0.53 |
| 1:A:805:LEU:HD22 | 1:A:928:LEU:HD21 | 1.90 | 0.53 |

*Continued on next page...*

For Manuscript Review

*Continued from previous page...*

| Atom-1 | Atom-2 | Interatomic distance (Å) | Clash overlap (Å) |
|---|---|---|---|
| 1:B:425:ASN:OD1 | 1:B:427:ASN:N | 2.39 | 0.53 |
| 1:A:1011:LYS:NZ | 1:A:1025:PHE:O | 2.42 | 0.52 |
| 1:B:305:ASN:O | 1:B:305:ASN:ND2 | 2.42 | 0.52 |
| 1:B:1118:ASN:OD1 | 1:B:1119:THR:N | 2.42 | 0.52 |
| 1:C:298:LYS:HG3 | 1:C:651:ILE:HD11 | 1.92 | 0.52 |
| 1:B:189:ARG:O | 1:B:195:LEU:HD12 | 2.10 | 0.52 |
| 1:A:312:LYS:HD2 | 1:A:521:ILE:HD13 | 1.91 | 0.52 |
| 1:C:750:LEU:HD21 | 1:C:991:VAL:HG22 | 1.91 | 0.52 |
| 1:A:1060:THR:HG22 | 1:A:1078:PHE:O | 2.09 | 0.52 |
| 1:A:539:LEU:HD23 | 1:A:574:ILE:HG22 | 1.92 | 0.51 |
| 1:A:356:LEU:HD23 | 1:A:362:PHE:CE2 | 2.46 | 0.51 |
| 1:A:286:GLU:OE1 | 1:A:616:LEU:HD21 | 2.11 | 0.51 |
| 1:A:967:LEU:HD13 | 1:A:971:GLU:OE1 | 2.11 | 0.51 |
| 1:C:809:VAL:HG23 | 1:C:928:LEU:HD12 | 1.93 | 0.51 |
| 1:A:1110:ASP:OD1 | 1:A:1110:ASP:O | 2.29 | 0.51 |
| 1:A:187:VAL:HG23 | 1:A:217:LEU:HD23 | 1.92 | 0.51 |
| 1:A:697:ILE:HD11 | 1:A:1058:PHE:CE2 | 2.46 | 0.51 |
| 1:A:187:VAL:HG23 | 1:A:217:LEU:CD2 | 2.40 | 0.51 |
| 1:C:560:THR:O | 1:C:574:ILE:HD11 | 2.11 | 0.50 |
| 1:C:352:ASP:O | 1:C:355:VAL:HG12 | 2.11 | 0.50 |
| 1:B:560:THR:O | 1:B:574:ILE:HD11 | 2.11 | 0.50 |
| 1:A:632:THR:HG23 | 1:A:657:ILE:HD12 | 1.95 | 0.49 |
| 1:C:627:ASN:C | 1:C:627:ASN:OD1 | 2.50 | 0.49 |
| 1:C:286:GLU:OE1 | 1:C:616:LEU:HD21 | 2.12 | 0.49 |
| 1:C:445:ARG:NE | 1:C:455:ASP:OD2 | 2.43 | 0.49 |
| 1:C:839:ASN:ND2 | 1:C:949:LEU:HD22 | 2.28 | 0.49 |
| 1:B:298:LYS:HG3 | 1:B:651:ILE:HD11 | 1.94 | 0.48 |
| 1:A:352:ASP:O | 1:A:355:VAL:HG12 | 2.13 | 0.48 |
| 1:B:682:LEU:HD21 | 1:C:852:MET:SD | 2.54 | 0.48 |
| 1:A:627:ASN:OD1 | 1:A:627:ASN:C | 2.52 | 0.48 |
| 1:C:39:ARG:NH2 | 1:C:185:GLU:OE2 | 2.39 | 0.48 |
| 1:C:632:THR:HG23 | 1:C:657:ILE:HD12 | 1.96 | 0.48 |
| 1:A:356:LEU:HD23 | 1:A:362:PHE:HE2 | 1.78 | 0.48 |
| 1:A:745:GLN:NE2 | 1:C:989:THR:HG21 | 2.29 | 0.48 |
| 1:A:928:LEU:HD22 | 1:A:931:LEU:HD12 | 1.96 | 0.47 |
| 1:A:722:ASN:O | 1:A:727:GLY:N | 2.36 | 0.47 |
| 1:B:539:LEU:HD23 | 1:B:574:ILE:HG22 | 1.95 | 0.47 |
| 1:B:722:ASN:O | 1:B:727:GLY:N | 2.37 | 0.47 |
| 1:C:129:CYS:HA | 1:C:160:CYS:CB | 2.44 | 0.47 |
| 1:A:745:GLN:HE22 | 1:C:989:THR:HG21 | 1.79 | 0.47 |
| 1:A:719:VAL:HG11 | 1:A:987:LEU:HD21 | 1.97 | 0.47 |

*Continued on next page...*

For Manuscript Review

Continued from previous page...

| Atom-1 | Atom-2 | Interatomic distance (Å) | Clash overlap (Å) |
|---|---|---|---|
| 1:B:261:THR:HG22 | 1:B:262:THR:N | 2.30 | 0.47 |
| 1:B:446:ARG:NH2 | 1:B:462:SER:O | 2.46 | 0.47 |
| 1:B:839:ASN:ND2 | 1:B:949:LEU:HD22 | 2.30 | 0.47 |
| 1:B:607:VAL:N | 1:B:608:PRO:HD2 | 2.30 | 0.46 |
| 1:A:333:THR:HG23 | 1:A:334:THR:CG2 | 2.43 | 0.46 |
| 1:C:187:VAL:HG23 | 1:C:217:LEU:CD2 | 2.45 | 0.46 |
| 1:A:261:THR:HG22 | 1:A:262:THR:N | 2.31 | 0.46 |
| 1:B:695:ILE:HD12 | 1:C:879:ILE:HG13 | 1.97 | 0.46 |
| 1:A:42:TYR:CD2 | 1:A:187:VAL:HG11 | 2.51 | 0.46 |
| 1:C:128:ALA:O | 1:C:160:CYS:HB2 | 2.16 | 0.46 |
| 1:B:1055:GLU:CG | 1:C:877:LEU:HD22 | 2.45 | 0.46 |
| 1:C:799:SER:OG | 1:C:802:GLU:OE1 | 2.28 | 0.46 |
| 1:B:968:ASP:N | 1:B:968:ASP:OD1 | 2.49 | 0.45 |
| 1:A:291:VAL:HG12 | 1:A:291:VAL:O | 2.15 | 0.45 |
| 1:B:187:VAL:HG23 | 1:B:217:LEU:HD23 | 1.98 | 0.45 |
| 1:C:968:ASP:OD1 | 1:C:968:ASP:N | 2.48 | 0.45 |
| 1:B:291:VAL:HG12 | 1:B:291:VAL:O | 2.17 | 0.45 |
| 1:B:627:ASN:OD1 | 1:B:627:ASN:C | 2.55 | 0.45 |
| 1:C:662:HIS:HB2 | 1:C:676:VAL:HG21 | 1.99 | 0.45 |
| 1:A:330:PHE:CD2 | 1:A:356:LEU:HD21 | 2.52 | 0.45 |
| 1:B:398:ILE:HD11 | 1:B:406:ILE:HG21 | 1.99 | 0.45 |
| 1:A:968:ASP:OD1 | 1:A:968:ASP:N | 2.49 | 0.45 |
| 1:B:688:ILE:HG23 | 1:B:688:ILE:O | 2.17 | 0.45 |
| 1:C:330:PHE:CD2 | 1:C:356:LEU:HD21 | 2.52 | 0.45 |
| 1:C:356:LEU:HD23 | 1:C:362:PHE:CE2 | 2.52 | 0.45 |
| 1:A:607:VAL:N | 1:A:608:PRO:HD2 | 2.31 | 0.45 |
| 1:A:877:LEU:HD22 | 1:C:1055:GLU:HG2 | 1.98 | 0.45 |
| 1:A:1075:GLU:N | 1:A:1075:GLU:OE1 | 2.50 | 0.45 |
| 1:B:656:GLY:N | 1:C:847:LEU:O | 2.48 | 0.44 |
| 1:C:291:VAL:HG12 | 1:C:291:VAL:O | 2.16 | 0.44 |
| 1:C:631:GLN:NE2 | 1:C:635:GLY:O | 2.50 | 0.44 |
| 1:C:187:VAL:HG23 | 1:C:217:LEU:HD23 | 1.99 | 0.44 |
| 1:C:377:ASP:OD1 | 1:C:377:ASP:O | 2.36 | 0.44 |
| 1:A:377:ASP:OD1 | 1:A:377:ASP:O | 2.35 | 0.44 |
| 1:A:549:PHE:CD1 | 1:A:549:PHE:C | 2.91 | 0.44 |
| 1:A:129:CYS:HA | 1:A:160:CYS:CB | 2.44 | 0.44 |
| 1:A:398:ILE:HD11 | 1:A:406:ILE:HG21 | 2.00 | 0.44 |
| 1:B:931:LEU:O | 1:B:934:VAL:HG12 | 2.18 | 0.44 |
| 1:C:607:VAL:N | 1:C:608:PRO:HD2 | 2.31 | 0.44 |
| 1:A:989:THR:HG21 | 1:B:745:GLN:NE2 | 2.33 | 0.44 |
| 1:B:802:GLU:O | 1:B:806:PHE:HD1 | 2.00 | 0.44 |

Continued on next page...

For Manuscript Review

*Continued from previous page...*

| Atom-1 | Atom-2 | Interatomic distance (Å) | Clash overlap (Å) |
|---|---|---|---|
| 1:C:313:GLU:HG2 | 1:C:315:VAL:HG23 | 1.99 | 0.44 |
| 1:B:445:ARG:NE | 1:B:455:ASP:OD2 | 2.49 | 0.44 |
| 1:C:109:ASN:O | 1:C:110:ASN:OD1 | 2.36 | 0.44 |
| 1:C:261:THR:HG22 | 1:C:262:THR:N | 2.33 | 0.44 |
| 1:B:187:VAL:HG23 | 1:B:217:LEU:CD2 | 2.47 | 0.43 |
| 1:C:542:SER:HB2 | 1:C:573:ASP:HB2 | 2.00 | 0.43 |
| 1:C:919:GLU:HA | 1:C:919:GLU:OE2 | 2.18 | 0.43 |
| 1:A:161:THR:HG21 | 1:C:454:ARG:HH12 | 1.82 | 0.43 |
| 1:A:931:LEU:O | 1:A:934:VAL:HG12 | 2.18 | 0.43 |
| 1:A:968:ASP:H | 1:C:371:SER:HG | 1.66 | 0.43 |
| 1:B:628:ASN:OD1 | 1:B:628:ASN:O | 2.36 | 0.43 |
| 1:A:370:VAL:HG21 | 1:A:375:LEU:HD21 | 2.01 | 0.43 |
| 1:C:582:VAL:HG22 | 1:C:599:TYR:CD1 | 2.53 | 0.43 |
| 1:B:305:ASN:ND2 | 1:B:305:ASN:C | 2.72 | 0.43 |
| 1:A:973:GLU:OE1 | 1:A:973:GLU:HA | 2.18 | 0.43 |
| 1:A:539:LEU:CD2 | 1:A:574:ILE:HG22 | 2.49 | 0.43 |
| 1:A:750:LEU:HD21 | 1:A:991:VAL:HG22 | 2.01 | 0.43 |
| 1:B:563:VAL:HG22 | 1:B:564:ARG:N | 2.34 | 0.43 |
| 1:B:697:ILE:HD11 | 1:B:1058:PHE:CE2 | 2.54 | 0.43 |
| 1:A:568:THR:HG22 | 1:A:568:THR:O | 2.19 | 0.43 |
| 1:B:265:LEU:HD22 | 1:B:273:ILE:HD13 | 1.99 | 0.43 |
| 1:A:298:LYS:HG3 | 1:A:651:ILE:HD11 | 2.01 | 0.43 |
| 1:A:802:GLU:O | 1:A:806:PHE:HD2 | 2.00 | 0.43 |
| 1:C:788:ILE:HG22 | 1:C:801:ILE:HD12 | 2.01 | 0.43 |
| 1:A:445:ARG:NE | 1:A:455:ASP:OD2 | 2.48 | 0.42 |
| 1:C:931:LEU:O | 1:C:934:VAL:HG12 | 2.19 | 0.42 |
| 1:A:877:LEU:HD21 | 1:C:1055:GLU:OE2 | 2.18 | 0.42 |
| 1:B:973:GLU:HA | 1:B:973:GLU:OE1 | 2.19 | 0.42 |
| 1:A:109:ASN:O | 1:A:110:ASN:OD1 | 2.37 | 0.42 |
| 1:A:504:LEU:HD13 | 1:B:966:ARG:HD2 | 2.01 | 0.42 |
| 1:A:717:THR:HG21 | 1:A:990:TYR:OH | 2.19 | 0.42 |
| 1:A:870:THR:HG21 | 1:A:877:LEU:HG | 2.01 | 0.42 |
| 1:A:685:ASP:OD1 | 1:A:686:SER:N | 2.52 | 0.42 |
| 1:B:109:ASN:O | 1:B:110:ASN:OD1 | 2.36 | 0.42 |
| 1:B:685:ASP:OD1 | 1:B:686:SER:N | 2.52 | 0.42 |
| 1:C:350:VAL:HG13 | 1:C:350:VAL:O | 2.20 | 0.42 |
| 1:C:688:ILE:O | 1:C:688:ILE:HG23 | 2.19 | 0.42 |
| 1:B:989:THR:O | 1:B:993:GLN:HG2 | 2.19 | 0.42 |
| 1:B:989:THR:HG21 | 1:C:745:GLN:NE2 | 2.34 | 0.42 |
| 1:A:688:ILE:O | 1:A:688:ILE:HG23 | 2.19 | 0.42 |
| 1:C:973:GLU:OE1 | 1:C:973:GLU:HA | 2.19 | 0.42 |

*Continued on next page...*

For Manuscript Review

*Continued from previous page...*

| Atom-1 | Atom-2 | Interatomic distance (Å) | Clash overlap (Å) |
|---|---|---|---|
| 1:A:695:ILE:CG2 | 1:A:1060:THR:HG21 | 2.49 | 0.42 |
| 1:B:371:SER:HG | 1:C:968:ASP:H | 1.68 | 0.42 |
| 1:C:808:LYS:NZ | 1:C:924:THR:HA | 2.34 | 0.42 |
| 1:B:352:ASP:O | 1:B:355:VAL:HG12 | 2.20 | 0.42 |
| 1:B:960:LEU:HD22 | 1:B:976:ILE:CD1 | 2.50 | 0.42 |
| 1:A:332:ALA:O | 1:A:496:ARG:NH1 | 2.49 | 0.41 |
| 1:B:165:VAL:O | 1:B:165:VAL:HG12 | 2.20 | 0.41 |
| 1:B:632:THR:HG23 | 1:B:657:ILE:HD12 | 2.01 | 0.41 |
| 1:A:1101:ASP:OD1 | 1:A:1102:ASN:N | 2.53 | 0.41 |
| 1:C:116:ILE:O | 1:C:116:ILE:HG23 | 2.20 | 0.41 |
| 1:A:443:TRP:CE3 | 1:A:444:VAL:HG13 | 2.55 | 0.41 |
| 1:A:615:GLN:HB2 | 1:A:623:TYR:HD1 | 1.86 | 0.41 |
| 1:A:624:SER:O | 1:A:625:THR:OG1 | 2.39 | 0.41 |
| 1:B:799:SER:OG | 1:B:802:GLU:OE1 | 2.34 | 0.41 |
| 1:B:1101:ASP:OD1 | 1:B:1102:ASN:N | 2.53 | 0.41 |
| 1:A:1073:PRO:HB3 | 1:A:1087:ILE:HD11 | 2.03 | 0.41 |
| 1:B:913:ALA:O | 1:B:917:ILE:HG22 | 2.21 | 0.41 |
| 1:B:539:LEU:CD2 | 1:B:574:ILE:HG22 | 2.51 | 0.41 |
| 1:C:578:SER:O | 1:C:579:PHE:HB3 | 2.21 | 0.41 |
| 1:C:800:PHE:CE2 | 1:C:918:GLN:NE2 | 2.85 | 0.41 |
| 1:C:885:MET:CE | 1:C:1032:LEU:HD13 | 2.51 | 0.41 |
| 1:A:839:ASN:ND2 | 1:A:949:LEU:HD22 | 2.35 | 0.41 |
| 1:C:275:ASP:O | 1:C:275:ASP:CG | 2.59 | 0.41 |
| 1:C:568:THR:O | 1:C:568:THR:HG22 | 2.20 | 0.41 |
| 1:A:459:ASP:OD1 | 1:A:459:ASP:O | 2.39 | 0.41 |
| 1:B:568:THR:O | 1:B:568:THR:HG22 | 2.21 | 0.41 |
| 1:B:870:THR:HG21 | 1:B:877:LEU:HG | 2.03 | 0.41 |
| 1:C:913:ALA:O | 1:C:917:ILE:HG22 | 2.21 | 0.41 |
| 1:A:425:ASN:OD1 | 1:A:426:THR:N | 2.54 | 0.41 |
| 1:A:1055:GLU:HG2 | 1:B:877:LEU:HD22 | 2.02 | 0.41 |
| 1:C:107:THR:OG1 | 1:C:112:SER:OG | 2.06 | 0.41 |
| 1:A:116:ILE:O | 1:A:116:ILE:HG23 | 2.20 | 0.40 |
| 1:A:615:GLN:HB2 | 1:A:623:TYR:CD1 | 2.56 | 0.40 |
| 1:B:356:LEU:HD23 | 1:B:362:PHE:CE2 | 2.56 | 0.40 |
| 1:B:518:THR:HG22 | 1:B:519:ASP:N | 2.37 | 0.40 |
| 1:C:976:ILE:HG22 | 1:C:980:ILE:HD12 | 2.03 | 0.40 |
| 1:B:923:THR:O | 1:B:924:THR:C | 2.59 | 0.40 |
| 1:C:1101:ASP:OD1 | 1:C:1102:ASN:N | 2.55 | 0.40 |

There are no symmetry-related clashes.

For Manuscript Review

#### 5.3 Torsion angles (i)

##### 5.3.1 Protein backbone (i)

In the following table, the Percentiles column shows the percent Ramachandran outliers of the chain as a percentile score with respect to all PDB entries followed by that with respect to all EM entries.

The Analysed column shows the number of residues for which the backbone conformation was analysed, and the total number of residues.

| Mol | Chain | Analysed | Favoured | Allowed | Outliers | Percentiles | |
|---|---|---|---|---|---|---|---|
| 1 | A | 947/1271 (74%) | 907 (96%) | 40 (4%) | 0 | 100 | 100 |
| 1 | B | 946/1271 (74%) | 911 (96%) | 35 (4%) | 0 | 100 | 100 |
| 1 | C | 948/1271 (75%) | 909 (96%) | 39 (4%) | 0 | 100 | 100 |
| All | All | 2841/3813 (74%) | 2727 (96%) | 114 (4%) | 0 | 100 | 100 |

There are no Ramachandran outliers to report.

##### 5.3.2 Protein sidechains (i)

In the following table, the Percentiles column shows the percent sidechain outliers of the chain as a percentile score with respect to all PDB entries followed by that with respect to all EM entries.

The Analysed column shows the number of residues for which the sidechain conformation was analysed, and the total number of residues.

| Mol | Chain | Analysed | Rotameric | Outliers | Percentiles | |
|---|---|---|---|---|---|---|
| 1 | A | 846/1099 (77%) | 829 (98%) | 17 (2%) | 55 | 80 |
| 1 | B | 846/1099 (77%) | 834 (99%) | 12 (1%) | 67 | 86 |
| 1 | C | 847/1099 (77%) | 830 (98%) | 17 (2%) | 55 | 80 |
| All | All | 2539/3297 (77%) | 2493 (98%) | 46 (2%) | 61 | 82 |

All (46) residues with a non-rotameric sidechain are listed below:

| Mol | Chain | Res | Type |
|---|---|---|---|
| 1 | A | 275 | ASP |
| 1 | A | 319 | ASN |
| 1 | A | 322 | ASN |
| 1 | A | 335 | PHE |
| 1 | A | 365 | PHE |

*Continued on next page...*

For Manuscript Review

*Continued from previous page...*

| Mol | Chain | Res | Type |
|---|---|---|---|
| 1 | A | 377 | ASP |
| 1 | A | 446 | ARG |
| 1 | A | 542 | SER |
| 1 | A | 549 | PHE |
| 1 | A | 624 | SER |
| 1 | A | 763 | GLU |
| 1 | A | 808 | LYS |
| 1 | A | 920 | SER |
| 1 | A | 922 | THR |
| 1 | A | 924 | THR |
| 1 | A | 1050 | TYR |
| 1 | A | 1069 | LYS |
| 1 | B | 280 | SER |
| 1 | B | 305 | ASN |
| 1 | B | 319 | ASN |
| 1 | B | 335 | PHE |
| 1 | B | 365 | PHE |
| 1 | B | 446 | ARG |
| 1 | B | 516 | LEU |
| 1 | B | 600 | GLN |
| 1 | B | 624 | SER |
| 1 | B | 763 | GLU |
| 1 | B | 808 | LYS |
| 1 | B | 922 | THR |
| 1 | C | 50 | SER |
| 1 | C | 275 | ASP |
| 1 | C | 319 | ASN |
| 1 | C | 322 | ASN |
| 1 | C | 348 | ASN |
| 1 | C | 365 | PHE |
| 1 | C | 371 | SER |
| 1 | C | 377 | ASP |
| 1 | C | 446 | ARG |
| 1 | C | 600 | GLN |
| 1 | C | 624 | SER |
| 1 | C | 698 | PRO |
| 1 | C | 723 | MET |
| 1 | C | 763 | GLU |
| 1 | C | 878 | GLN |
| 1 | C | 922 | THR |
| 1 | C | 1069 | LYS |

Sometimes sidechains can be flipped to improve hydrogen bonding and reduce clashes. All (3) such

For Manuscript Review

sidechains are listed below:

| Mol | Chain | Res | Type |
|---|---|---|---|
| 1 | B | 159 | ASN |
| 1 | B | 628 | ASN |
| 1 | B | 896 | GLN |

###### 5.3.3 RNA (i)

There are no RNA molecules in this entry.

##### 5.4 Non-standard residues in protein, DNA, RNA chains (i)

There are no non-standard protein/DNA/RNA residues in this entry.

##### 5.5 Carbohydrates (i)

There are no monosaccharides in this entry.

##### 5.6 Ligand geometry (i)

There are no ligands in this entry.

##### 5.7 Other polymers (i)

There are no such residues in this entry.

##### 5.8 Polymer linkage issues (i)

There are no chain breaks in this entry.

For Manuscript Review

### 6 Map visualisation (i)

This section contains visualisations of the EMDB entry EMD-45402. These allow visual inspection of the internal detail of the map and identification of artifacts.

Images derived from a raw map, generated by summing the deposited half-maps, are presented below the corresponding image components of the primary map to allow further visual inspection and comparison with those of the primary map.

#### 6.1 Orthogonal projections (i)

##### 6.1.1 Primary map

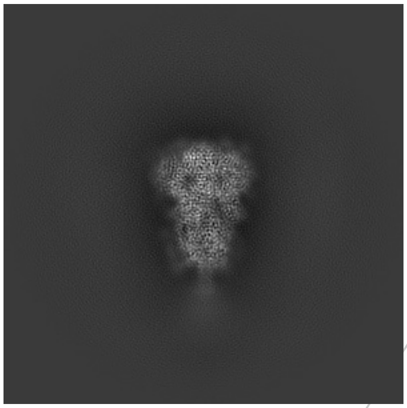
X

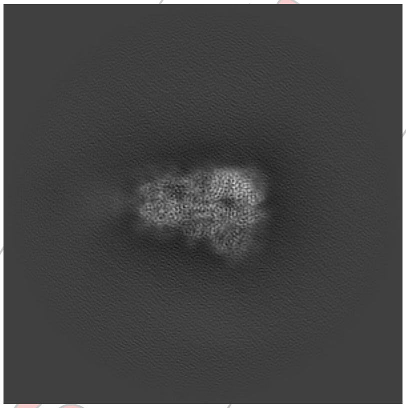
Y

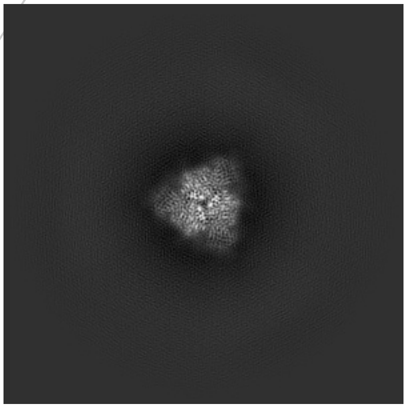
Z

##### 6.1.2 Raw map

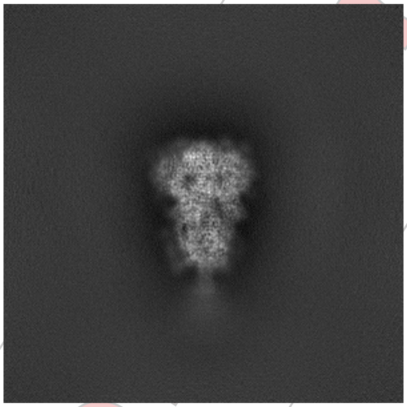
X

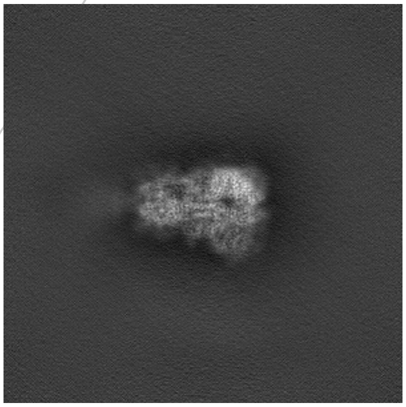
Y

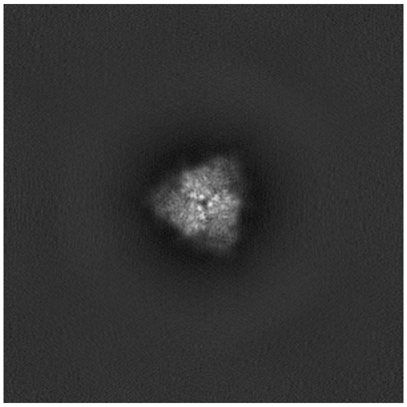
Z

The images above show the map projected in three orthogonal directions.

For Manuscript Review

#### 6.2 Central slices (i)

##### 6.2.1 Primary map

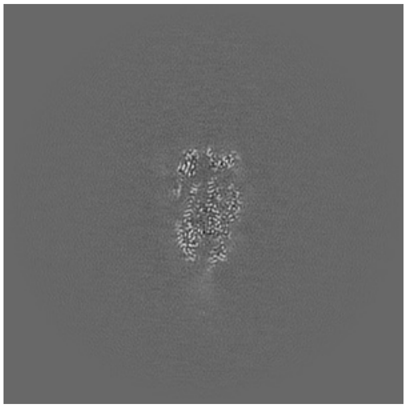

X Index: 256

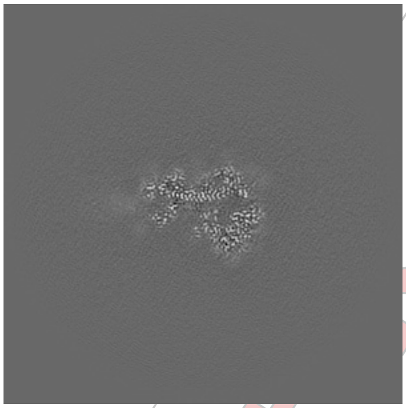

Y Index: 256

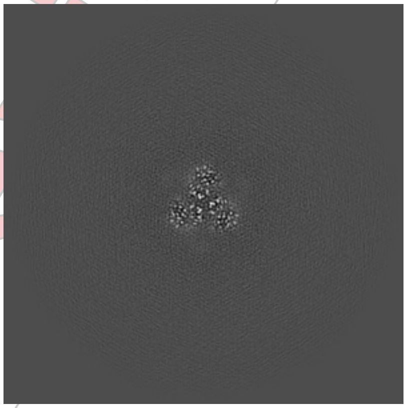

Z Index: 256

##### 6.2.2 Raw map

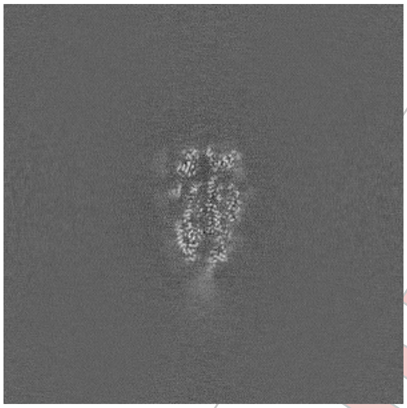

X Index: 256

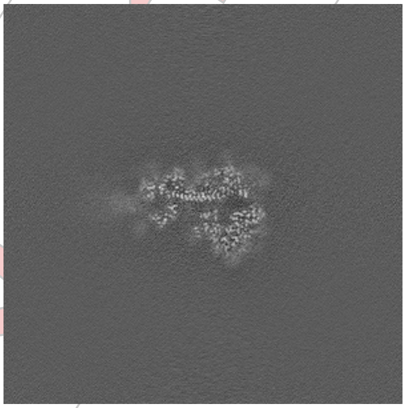

Y Index: 256

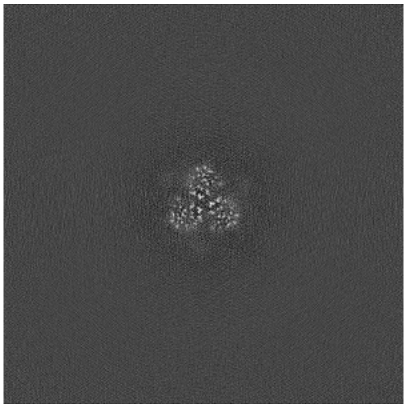

Z Index: 256

The images above show central slices of the map in three orthogonal directions.

For Manuscript Review

#### 6.3 Largest variance slices (i)

##### 6.3.1 Primary map

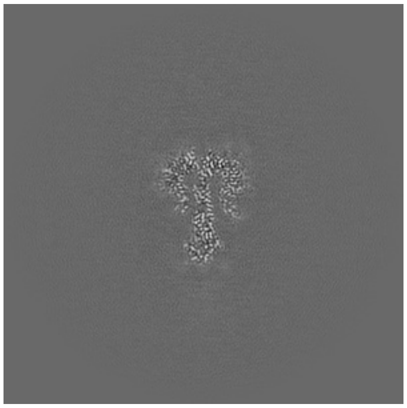

X Index: 272

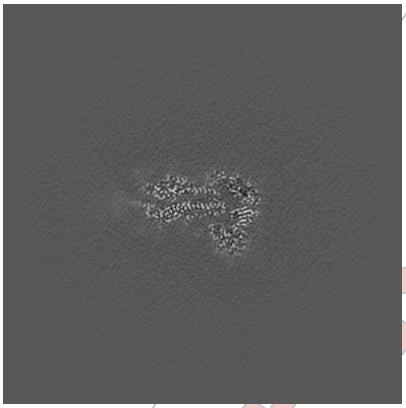

Y Index: 264

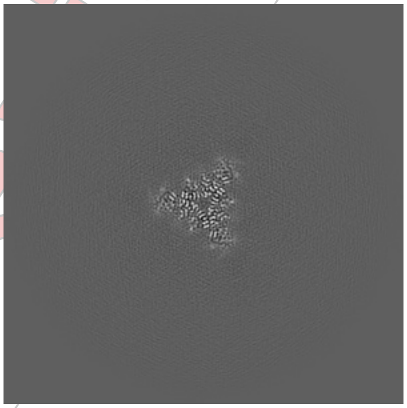

Z Index: 305

##### 6.3.2 Raw map

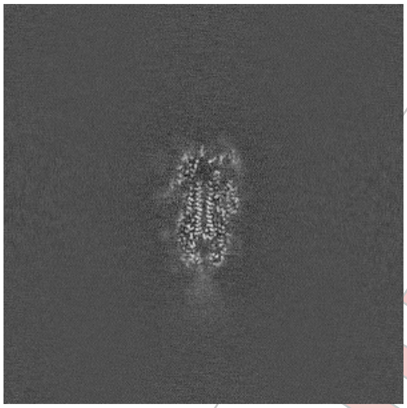

X Index: 251

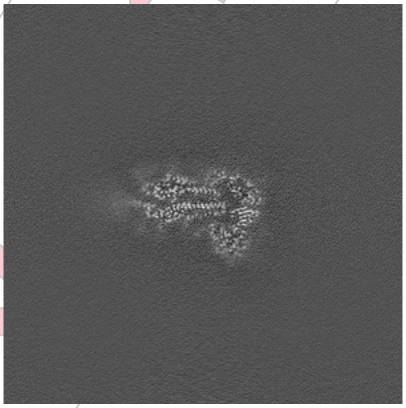

Y Index: 264

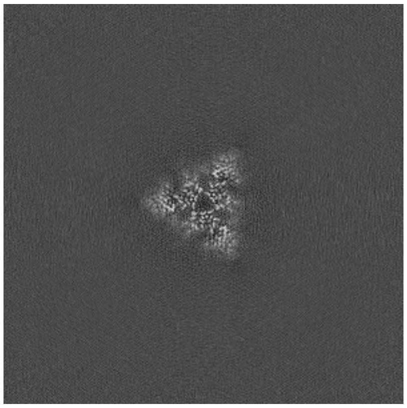

Z Index: 296

The images above show the largest variance slices of the map in three orthogonal directions.

For Manuscript Review

#### 6.4 Orthogonal standard-deviation projections (False-color) (i)

##### 6.4.1 Primary map

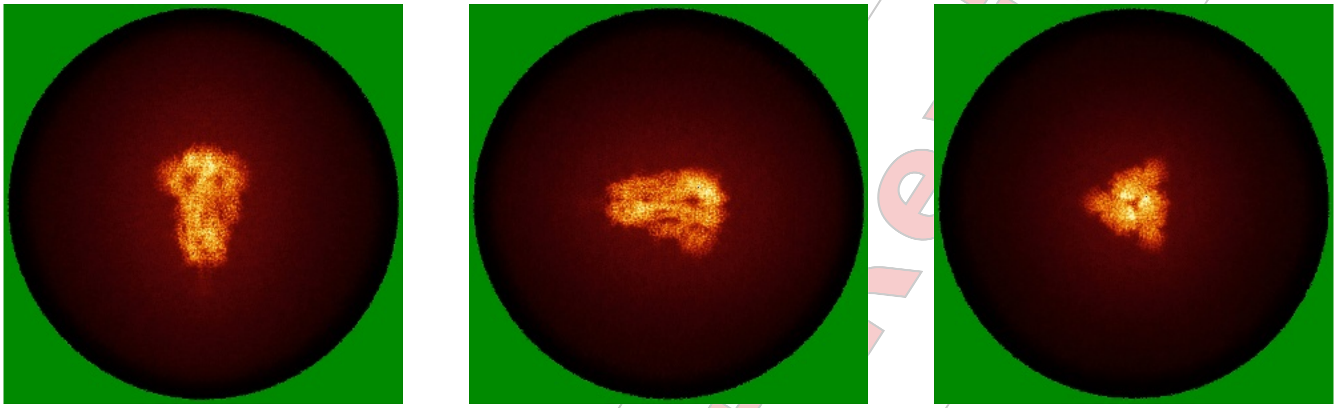

X Y Z

##### 6.4.2 Raw map

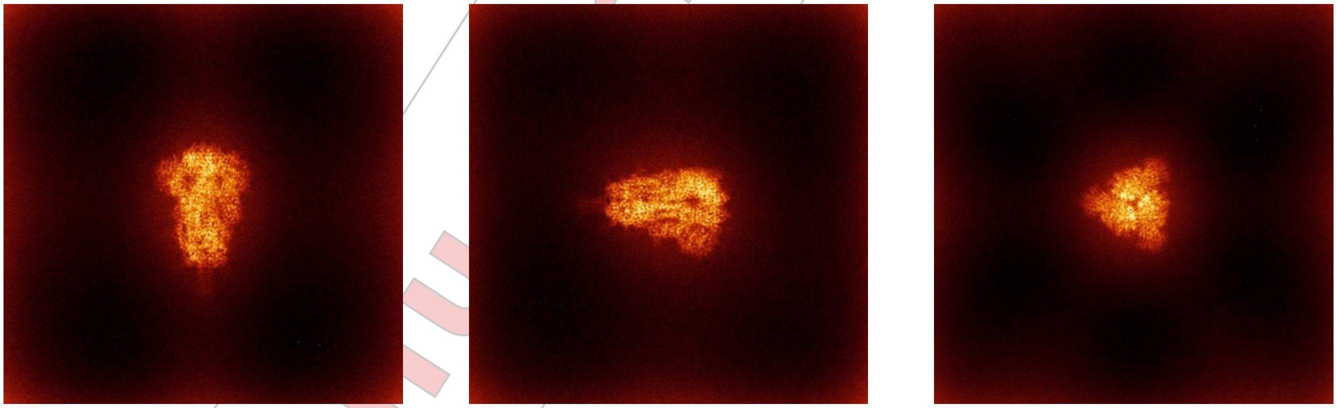

X Y Z

The images above show the map standard deviation projections with false color in three orthogonal directions. Minimum values are shown in green, max in blue, and dark to light orange shades represent small to large values respectively.

For Manuscript Review

#### 6.5 Orthogonal surface views (i)

##### 6.5.1 Primary map

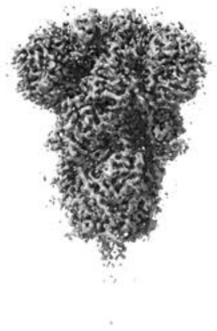

X

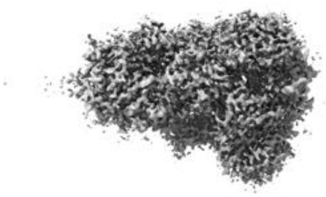

Y

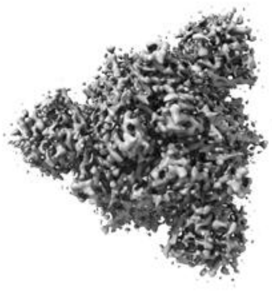

Z

The images above show the 3D surface view of the map at the recommended contour level 0.18. These images, in conjunction with the slice images, may facilitate assessment of whether an appropriate contour level has been provided.

##### 6.5.2 Raw map

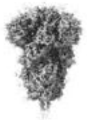

X

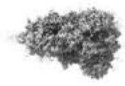

Y

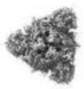

Z

These images show the 3D surface of the raw map. The raw map's contour level was selected so that its surface encloses the same volume as the primary map does at its recommended contour level.

#### 6.6 Mask visualisation (i)

This section was not generated. No masks/segmentation were deposited.

For Manuscript Review

### 7 Map analysis (i)

This section contains the results of statistical analysis of the map.

#### 7.1 Map-value distribution (i)

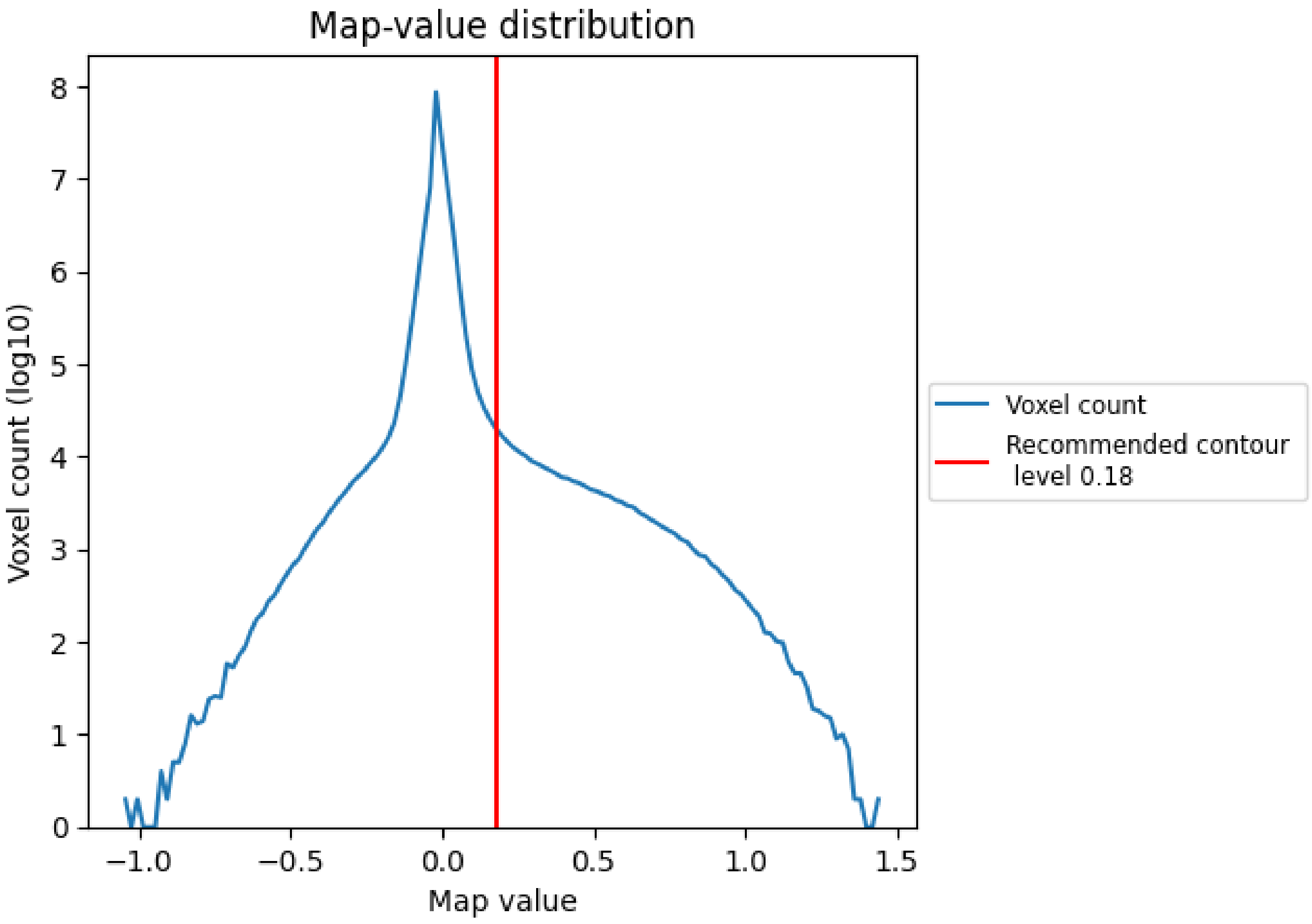


The map-value distribution is plotted in 128 intervals along the x-axis. The y-axis is logarithmic. A spike in this graph at zero usually indicates that the volume has been masked.

#### 7.2 Volume estimate (i)

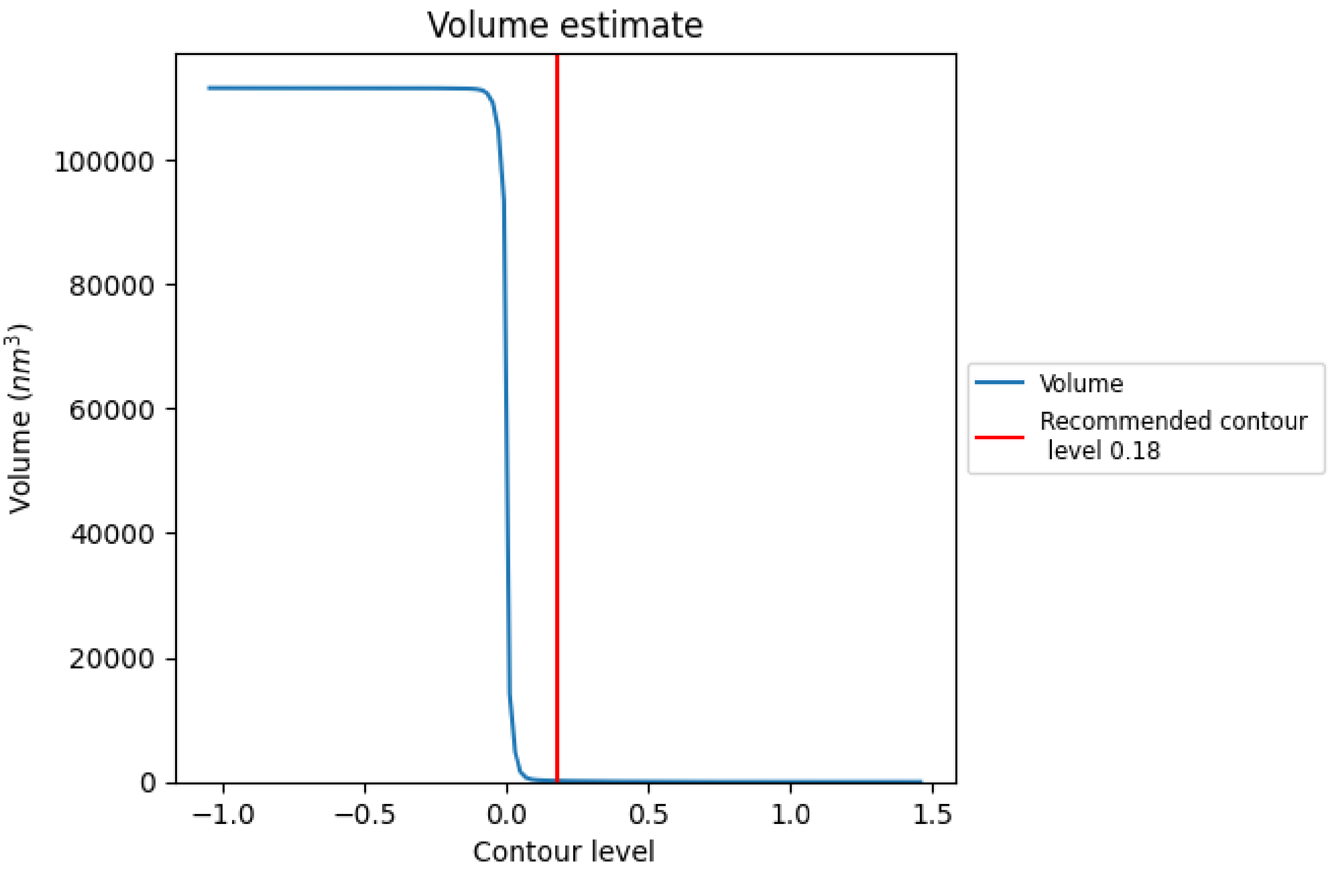


The volume at the recommended contour level is 167 nm$^3$; this corresponds to an approximate mass of 151 kDa.

The volume estimate graph shows how the enclosed volume varies with the contour level. The recommended contour level is shown as a vertical line and the intersection between the line and the curve gives the volume of the enclosed surface at the given level.

For Manuscript Review

#### 7.3 Rotationally averaged power spectrum (i)

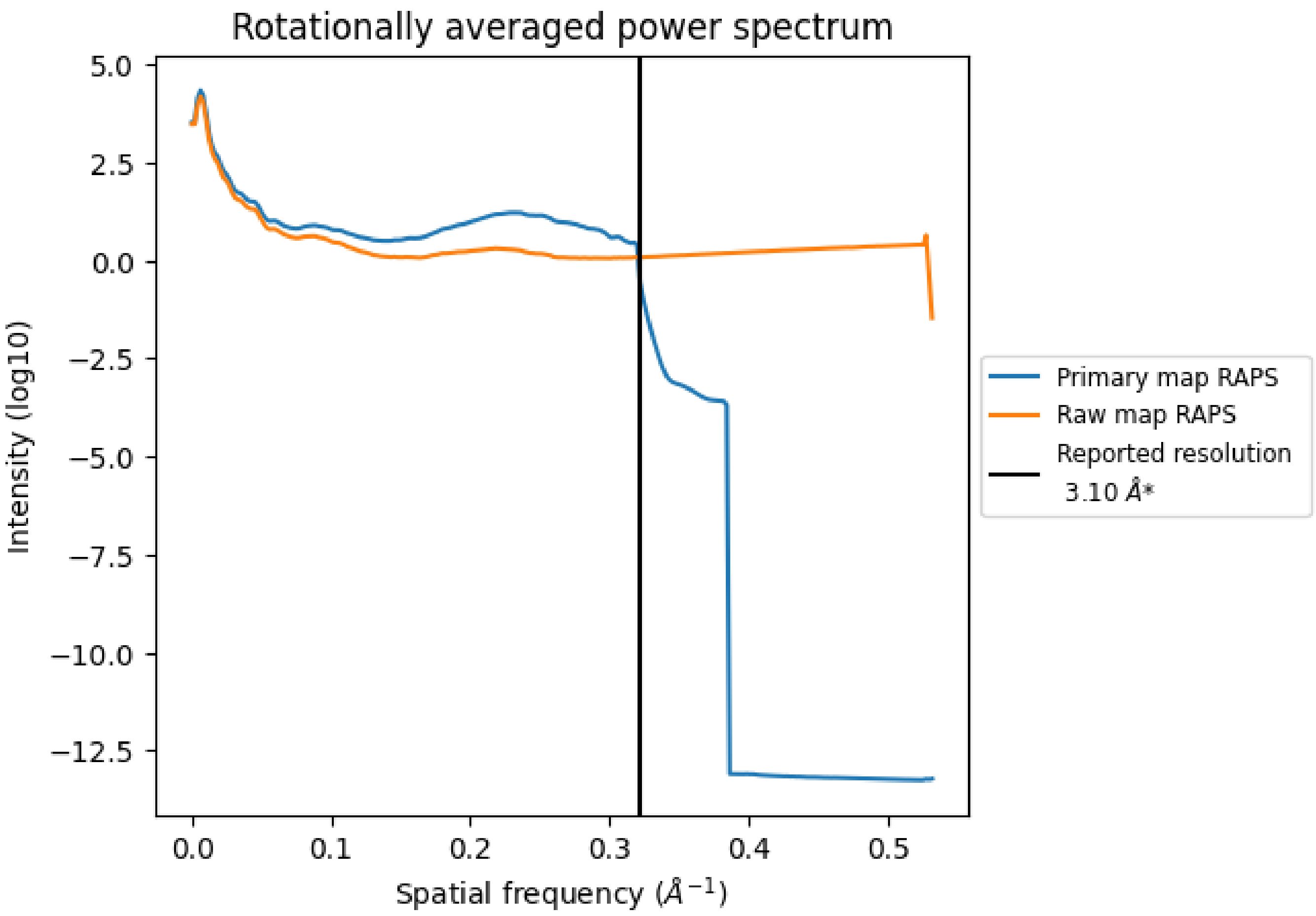


*Reported resolution corresponds to spatial frequency of 0.323 Å$^{-1}$

For Manuscript Review

### 8 Fourier-Shell correlation ⓘ

Fourier-Shell Correlation (FSC) is the most commonly used method to estimate the resolution of single-particle and subtomogram-averaged maps. The shape of the curve depends on the imposed symmetry, mask and whether or not the two 3D reconstructions used were processed from a common reference. The reported resolution is shown as a black line. A curve is displayed for the half-bit criterion in addition to lines showing the 0.143 gold standard cut-off and 0.5 cut-off.

#### 8.1 FSC ⓘ

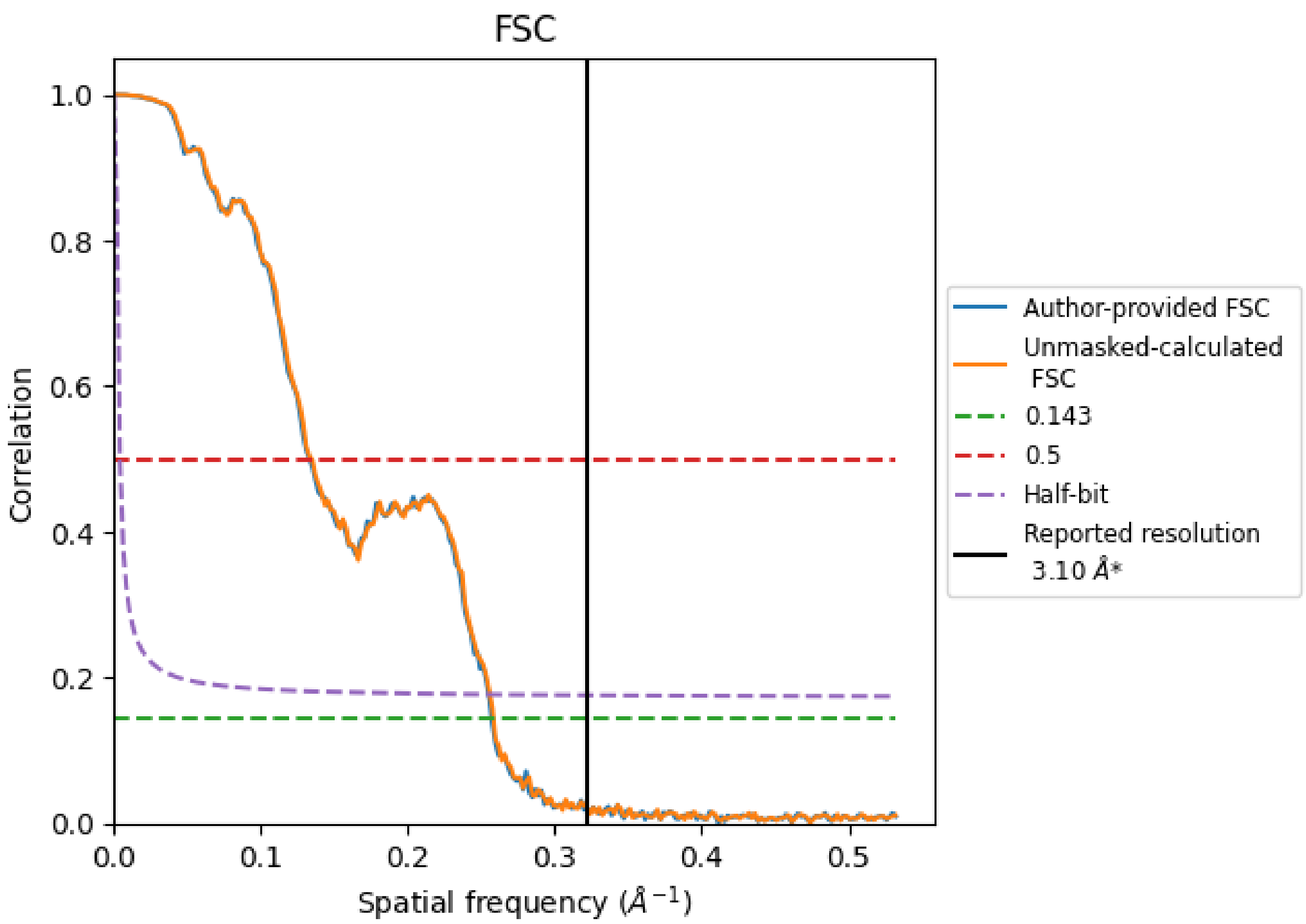


*Reported resolution corresponds to spatial frequency of 0.323 Å$^{-1}$

For Manuscript Review

#### 8.2 Resolution estimates (i)

| Resolution estimate (Å) | Estimation criterion (FSC cut-off) | | |
|---|---|---|---|
| | 0.143 | 0.5 | Half-bit |
| Reported by author | 3.10 | - | - |
| Author-provided FSC curve | 3.89 | 7.51 | 3.92 |
| Unmasked-calculated* | 3.87 | 7.45 | 3.91 |

*Resolution estimate based on FSC curve calculated by comparison of deposited half-maps. The value from author-provided FSC intersecting FSC 0.143 CUT-OFF 3.89 differs from the reported value 3.1 by more than 10 %

The value from deposited half-maps intersecting FSC 0.143 CUT-OFF 3.87 differs from the reported value 3.1 by more than 10 %

For Manuscript Review

### 9 Map-model fit (i)

This section contains information regarding the fit between EMDB map EMD-45402 and PDB model 9CAS. Per-residue inclusion information can be found in section 3 on page 10.

#### 9.1 Map-model overlay (i)

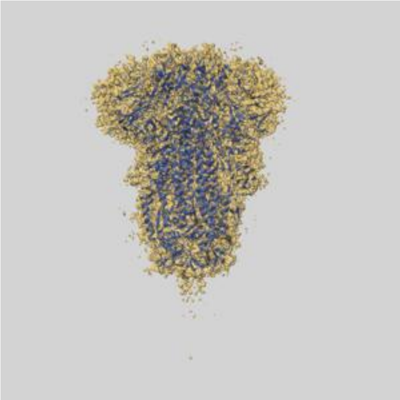

X

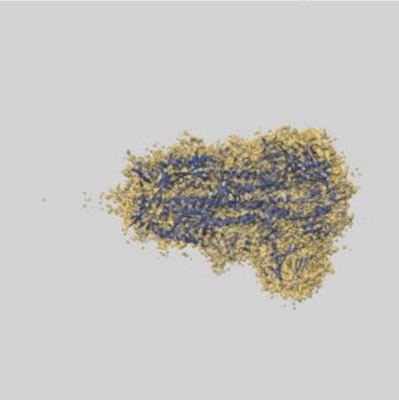

Y

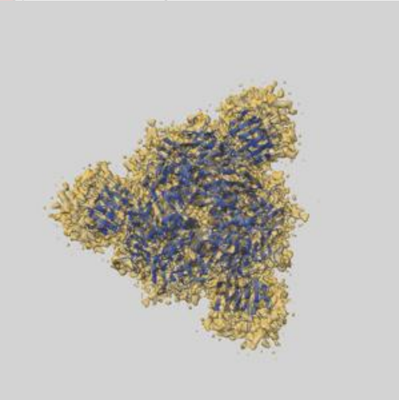

Z

The images above show the 3D surface view of the map at the recommended contour level 0.18 at 50% transparency in yellow overlaid with a ribbon representation of the model coloured in blue. These images allow for the visual assessment of the quality of fit between the atomic model and the map.

For Manuscript Review

#### 9.2 Q-score mapped to coordinate model (i)

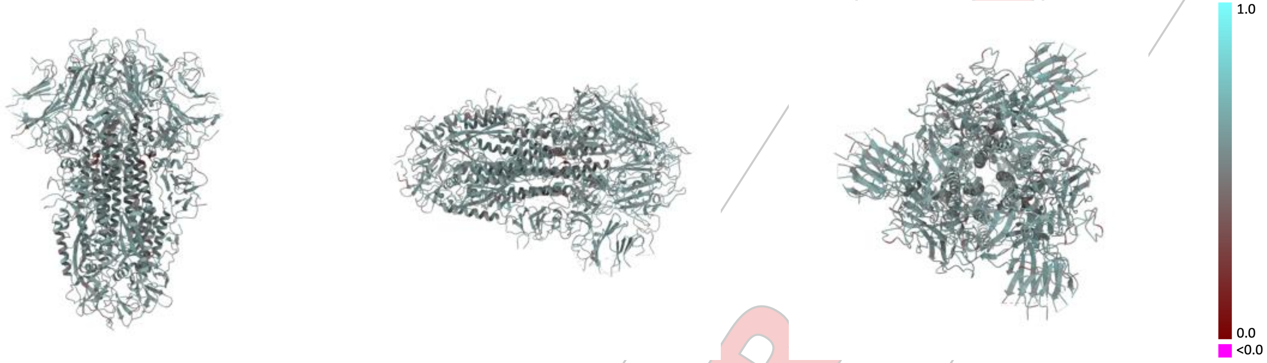


X Y Z

The images above show the model with each residue coloured according its Q-score. This shows their resolvability in the map with higher Q-score values reflecting better resolvability. Please note: Q-score is calculating the resolvability of atoms, and thus high values are only expected at resolutions at which atoms can be resolved. Low Q-score values may therefore be expected for many entries.

#### 9.3 Atom inclusion mapped to coordinate model (i)

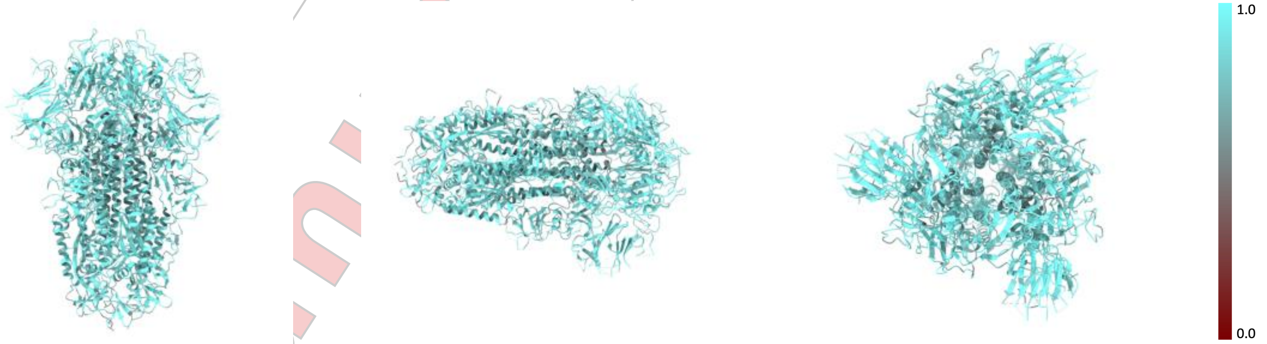


X Y Z

The images above show the model with each residue coloured according to its atom inclusion. This shows to what extent they are inside the map at the recommended contour level (0.18).

For Manuscript Review

#### 9.4 Atom inclusion (i)

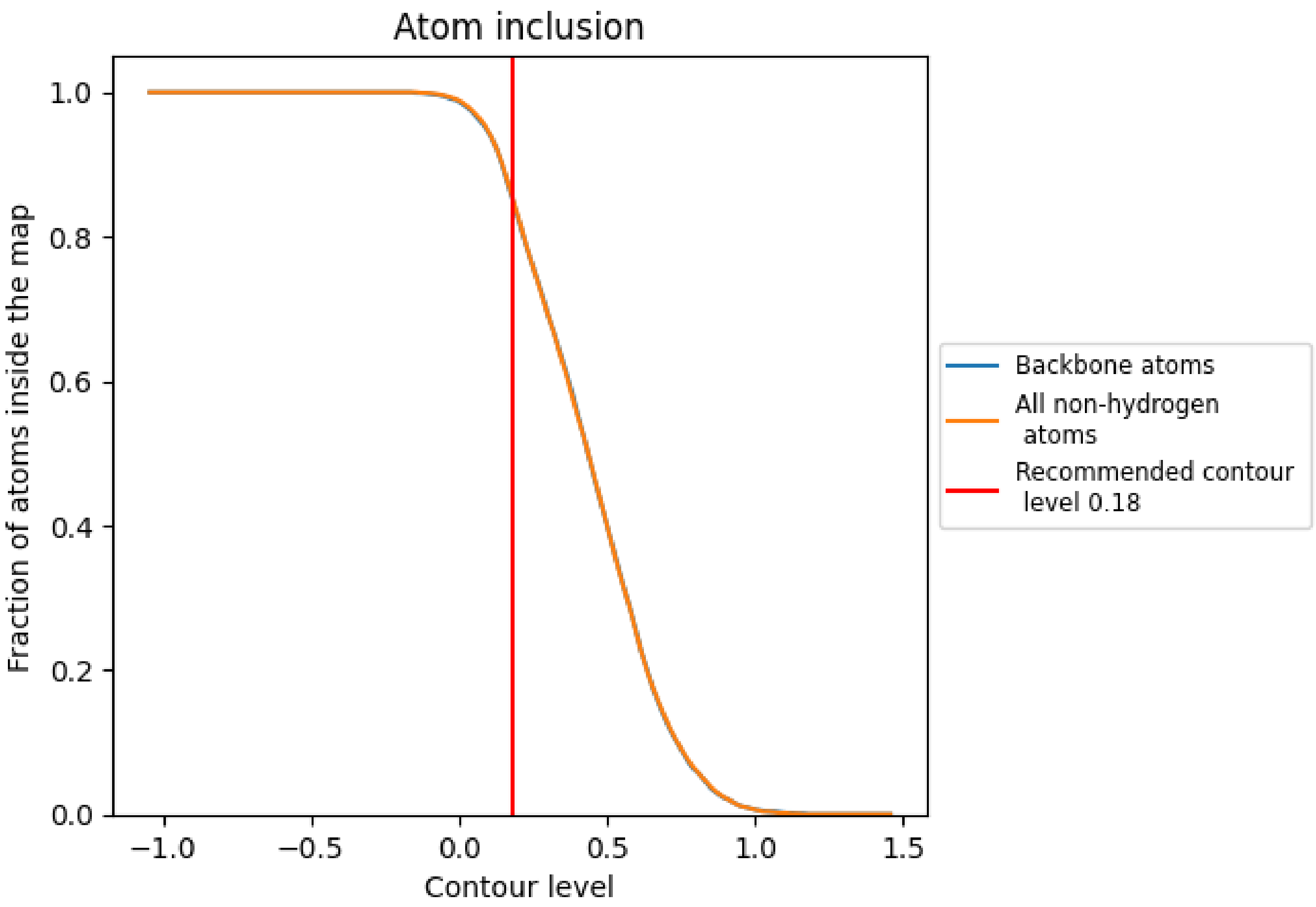


At the recommended contour level, 85% of all backbone atoms, 86% of all non-hydrogen atoms, are inside the map.

For Manuscript Review

#### 9.5 Map-model fit summary (i)

The table lists the average atom inclusion at the recommended contour level (0.18) and Q-score for the entire model and for each chain.

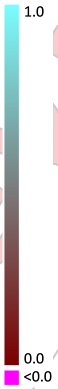


| Chain | Atom inclusion | Q-score |
|---|---|---|
| All | 0.8560 | 0.5470 |
| A | 0.8610 | 0.5470 |
| B | 0.8600 | 0.5470 |
| C | 0.8600 | 0.5470 |

For Manuscript Review
